# The AMIGO1 adhesion protein activates Kv2.1 voltage sensors

**DOI:** 10.1101/2021.06.20.448455

**Authors:** R.J. Sepela, R.G. Stewart, L.A. Valencia, P. Thapa, Z. Wang, B.E. Cohen, J.T. Sack

## Abstract

Kv2 voltage-gated potassium channels are modulated by AMIGO neuronal adhesion proteins. Here, we identify steps in the conductance activation pathway of Kv2.1 channels that are modulated by AMIGO1 using voltage clamp recordings and spectroscopy of heterologously expressed Kv2.1 and AMIGO1 in mammalian cell lines. AMIGO1 speeds early voltage sensor movements and shifts the gating charge–voltage relationship to more negative voltages. The gating charge–voltage relationship indicates that AMIGO1 exerts a larger energetic effect on voltage sensor movement than apparent from the midpoint of the conductance–voltage relationship. When voltage sensors are detained at rest by voltage sensor toxins, AMIGO1 has a greater impact on the conductance–voltage relationship. Fluorescence measurements from voltage sensor toxins bound to Kv2.1 indicate that with AMIGO1, the voltage sensors enter their earliest resting conformation, yet this conformation is less stable upon voltage stimulation. We conclude that AMIGO1 modulates the Kv2.1 conductance activation pathway by destabilizing the earliest resting state of the voltage sensors.

**Statement of Significance:** Kv2 potassium channels activate a potassium conductance that shapes neuronal action potentials. The AMIGO family of adhesion proteins modulate activation of Kv2 conductances, yet, which activation steps are modified is unknown. This study finds that AMIGO1 destabilizes the earliest resting conformation of the Kv2.1 voltage sensors to promote activation of channel conductance.

## Introduction

Voltage–gated potassium (Kv) channels of the Kv2 family open following membrane depolarization and are critical regulators of neuronal electrical excitability. Mammals have two Kv2 pore-forming α subunits, Kv2.1 and Kv2.2, which function as homo- or heterotetramers (1). The molecular architecture of Kv2 channels is similar to Kv1 channels for which atomic resolution structures have been solved (2). Each α subunit monomer has six transmembrane helical segments, S1-S6. S1-S4 comprise a voltage sensor domain (VSD) while S5 and S6 together form one quarter of the central pore domain. In response to sufficiently positive intracellular voltages, gating charges within the VSD translate from an intracellular resting position to a more extracellular activated conformation. This gating charge movement powers the conformational changes of voltage sensor activation, which are coupled to subsequent pore opening and K^+^ conduction (3). Kv channels progress through a landscape of conformations leading to opening, all of which define a pathway for the activation of the K^+^ conductance. The activation pathway of Kv2 channels is distinct from Kv1 channels, as Kv2.1 channels have a pore opening step which is slower and more weakly voltage-dependent than the VSD movement of Kv1 channels (3–5). The unique kinetics and voltage dependence of Kv2 currents are critical to neuronal activity, as they regulate action potential duration and can either support or limit repetitive firing (6–10).

Kv2 channels are abundant in most mammalian central neurons (11). Genetic deletion of Kv2.1 leads to seizure susceptibility and behavioral hyperexcitability in mice (12), and human Kv2.1 mutations result in developmental epileptic encephalopathy (13–15), underscoring the importance of these channels to brain function. Homeostatic Kv2.1 regulation maintains neuronal excitability (16). Kv2.1 regulation by ischemia (17, 18), glutamate (19), phosphorylation (20) and SUMOylation (21) and AMIGO auxiliary subunits (22, 23) all shift the midpoint of the conductance–voltage relation (*G–V*). However, it is not known which steps in the conductance activation pathway are modulated by any of these forms of regulation.

To identify steps in the Kv2.1 conduction activation pathway that are susceptible to modulation, we studied the impact of an AMIGO auxiliary subunit. The AMIGO (<u>AM</u>photerin–<u>I</u>nduced <u>G</u>ene and <u>O</u>pen reading frame) family of proteins contains three paralogs in mammals: AMIGO1, AMIGO2, and AMIGO3. AMIGO proteins are single-pass transmembrane proteins with an extracellular immunoglobulin domain and several leucine-rich repeats (24). AMIGO1 has been proposed to play a role in schizophrenia biology (25). In vertebrate brain neurons, AMIGO1 is important for cell adhesion (24), neuronal tract development (26), and circuit formation (25–27). AMIGO1 colocalizes with Kv2 in neurons throughout the brains of multiple mammalian species (22, 28). Coimmunoprecipitation of AMIGO1 and Kv2.1 (22, 23, 26) and co-diffusion through cell membranes (22) indicate a robust interaction, consistent with an AMIGO1–Kv2.1 complex being sufficiently stable for intensive biophysical studies. All three AMIGO proteins activate the conductance of both Kv2 channel subtypes, shifting the conductance–voltage relation by −5 to −15 mV (22, 23). While these shifts may seem small in excitable cells that can have voltage swings of more than 100 mV, human mutations that shift the conductance–voltage relation of ion channel gating by similar magnitudes are correlated with physiological consequences (13, 29–31). However, it is difficult to determine whether the physiological consequences of mutations are caused by the gating shifts themselves.

Here we investigate which steps in the Kv2.1 conductance activation pathway are modulated by AMIGO1. In other voltage-gated ion channels, the *G–V* relation can be shifted to more negative voltages by modulating pore opening (32–34), voltage sensor movement (35, 36), or voltage sensor-pore coupling (37–39). Single-pass transmembrane auxiliary subunits modulate other voltage-gated ion channel α subunits by a variety of mechanisms (32, 38, 40, 41). However, AMIGO1 only shares a limited degree of homology with other single-pass transmembrane auxiliary subunits (42), and divergent structural interactions have been observed among single-pass transmembrane auxiliary subunits (43, 44). As there is no consensus binding pose or mechanism of interaction for auxiliary subunits, it is difficult to predict on which step in the conductance activation pathway AMIGO1 acts. A recent study proposed that AMIGO proteins shift Kv2.1 conductance by increasing voltage sensor-pore coupling and that AMIGO-conferred changes to Kv2 voltage-sensing machinery are unlikely (23). Here we ask whether AMIGO1 alters conformational changes associated with pore opening or with voltage sensor movement using a combination of electrophysiological and imaging approaches. We find that AMIGO1 modulates voltage sensor movements which occur before pore opening. We find AMIGO1 to have a greater impact on early voltage sensor movements than the conductance– voltage relation. We conclude that AMIGO1 destabilizes the earliest resting conformation in the pathway of channel activation.

## Methods

### GxTX peptides

A conjugate of a cysteine–modified guangxitoxin–1E and the maleimide of fluorophore Alexa594 (GxTX Ser13Cys(Alexa594)) was used to selectively modulate Kv2.1 channel gating and to fluorescently identify surface-expressing Kv2.1 channels (45). Conjugates of propargylglycine (Pra)–modified GxTX and the fluorophore JP–N3 (GxTX Ser13Pra(JP) and GxTX Lys27Pra(JP)) were used to monitor the chemical environment surrounding GxTX when localized to the channel (46). All modified GxTX–mutants were synthesized by solid phase peptide synthesis as described (46–48). Stock solutions were stored at −80 °C and thawed on ice on the day of experiment.

### Cell culture and transfection

The HEK293 cell line subclone TS201A was a gift from Vladimir Yarov-Yarovoy and was maintained in DMEM (Gibco Cat# 11995-065) with 10% Fetal Bovine Serum (HyClone, SH30071.03HI, LotAXM55317) and 1% penicillin/streptomycin (Gibco, 15-140-122) in a humidified incubator at 37°C under 5% CO2. Chinese Hamster Ovary (CHO) cell lines were a Tetracycline-Regulated Expression (T-REx) variant (Invitrogen, Cat# R71807), and cultured as described previously (47). The Kv2.1–CHO cell subclone (49) was stably transfected with pCDNA4/TO encoding the rat Kv2.1 (rKv2.1) channel. Cell lines were negative for mycoplasma by biochemical test (Lonza, LT07). 1 *μ*g/ml minocycline (Enzo Life Sciences), prepared in 70% ethanol, was added to Kv2.1–CHO cells to induce rKv2.1 channel expression for 1.5 hours to minimize series resistance-induced voltage errors in K^+^ current recordings or for 48 hours to produce sufficient Kv2.1 density necessary for recording gating currents. 5 minutes prior to transfection, cells were plated at 40% confluency in unsupplemented culture media free of antibiotics, selection agents, and serum and allowed to settle at room temperature. For imaging studies (except concentration– response), cells were plated in 35 mm No. 1.5 glass–bottom dishes (MatTek, P35G-1.5-20-C). For concentration-response time–lapse imaging, cells were plated onto 22 x 22 mm No. 1.5H cover glass (Deckglaser). For electrophysiological studies, cells were plated in 35 mm tissue culture treated polystyrene dishes (Fisher Scientific, 12-556-000). Transfections were achieved with Lipofectamine 2000 (Life Technologies, 11668-027). Each transfection included 220 *μ*L Opti–MEM (Life Technologies, 31985062), 1.1 *μ*L Lipofectamine, and the following amount of plasmid DNA. HEK293 cell experiments: 0.1 *μ*g of mKv2.1 DNA and either 0.1 *μ*g of pEGFP, mAMIGO1–pIRES2–GFP DNA, or hSCN1β–pIRES2–GFP. The pIRES2–GFP vector has an encoded internal ribosome entry site which promotes continuous translation of two genes from a singular mRNA (50) so that GFP fluorescence indicates the presence of AMIGO1 or SCN1β mRNA. Kv2.1–CHO cell experiments: 1 *μ*g of either mAMIGO1–pEYFP–N1, pEGFP, rAMIGO2–pEYFP–N1, or rAMIGO3–pEYFP–N1. CHO cell experiments: 1 *μ*g of both pCAG–ChroME–mRuby2-ST and mAMIGO1-pEYFP-N1. Cells were incubated in the transfection cocktail and 2 mL of unsupplemented media for 6-8 hours before being returned to regular growth media, and used for experiments 40-48 hours after transfection. pEGFP, mAMIGO1–pEYFP–N1, and pCAG–ChroME-mRuby2-ST (51) plasmids were gifts from James Trimmer. mAMIGO1–pEYFP–N1 uses a VPRARDPPVAT linker to tag the internal C–terminus of wild–type mouse AMIGO1 (NM_001004293.2 or NM_146137.3) with eYFP. pCAG– ChroME–mRuby2–ST encodes an mRuby2–tagged channelrhodopsin with a Kv2.1 PRC trafficking sequence (51, 52). mKv2.1 (NM_008420) was purchased from OriGene (MG210968). hSCN1β–pIRES2–GFP was a gift from Vladimir Yarov-Yarovoy. mAMIGO1 was subcloned into pIRES2–GFP between NheI and BamHI restriction sites. rAMIGO2–pEYFP–N1 and rAMIGO3–pEYFP–N1 were generated by subcloning rat AMIGO2 (NM_182816.2) or rat AMIGO3 (NM_178144.1) in place of mAMIGO1 in the mAMIGO1–pEYFP–N1 vector.

### Whole-cell K^+^ ionic currents

Voltage clamp was achieved with an Axopatch 200B patch clamp amplifier (Axon Instruments) run by Patchmaster (HEKA). Solutions: HEK293 internal (in mM) 160 KCl, 5 EGTA, 10 HEPES, 1 CaCl_2_, 2 MgCl_2_, and 10 glucose, adjusted to pH 7.3 with KOH, 345 mOsm. HEK293 external (in mM) 5 KCl, 160 NaCl, 10 HEPES, 2 CaCl_2_, 2 MgCl_2_, 10 glucose, pH 7.3 with NaOH, 345 mOsm, 5 *μ*M tetrodotoxin added to recording solution: LJP 3.9 mV, E_K_: −89.0 mV with HEK293 internal. Kv2.1–CHO internal (in mM) 70 KCl, 5 EGTA, 50 HEPES, 50 KF, and 35 KOH, adjusted to pH 7.4 with KOH, 310 mOsm. Kv2.1–CHO external (in mM) 3.5 KCl, 155 NaCl, 10 HEPES, 1.5 CaCl_2_, 1 MgCl_2_, adjusted to pH 7.4 with NaOH, 315 mOsm: LJP 8.5 mV, E_K_: −97.4 mV with Kv2.1– CHO cell internal. High Mg^2+^ Kv2.1–CHO external (in mM) 3.5 KCl, 6.5 NaCl, 10 HEPES, 1.5 CaCl_2_, 100 MgCl_2_, adjusted to pH 7.4 with NaOH, 289 mOsm: LJP 13.1 mV, E_K_: −97.4 mV with Kv2.1–CHO internal. Osmolality measured with a vapor pressure osmometer (Wescor, 5520), 5% difference between batches were tolerated. Liquid junction potential (LJP) values were tabulated using Patcher’s Power Tools version 2.15 (Max-Planck), and corrected *post hoc*, during analysis. Voltage protocols list command voltages, prior to LJP correction. Kv2.1–CHO cells were harvested by scraping in Versene (Gibco, 15040066) or TrypLE (Gibco, 12563011). HEK293 cells were dislodged by scraping. Cells were washed three times in a polypropylene tube in the external solution used in the recording chamber bath by pelleting at 1,000 x *g* for 2 min, and rotated at room temperature (22-24 °C) until use. Cells were then pipetted into a 50 *μ*L recording chamber (Warner Instruments, RC-24N) and allowed to settle for 5 or more minutes. After adhering to the bottom of the glass recording chamber, cells were rinsed with external solution using a gravity–driven perfusion system. Cells showing plasma membrane-associated YFP, or intracellular GFP of intermediate intensity, were selected for patching. Thin-wall borosilicate glass recording pipettes (BF150-110-7.5HP, Sutter) were pulled with blunt tips, coated with silicone elastomer (Sylgard 184, Dow Corning), heat cured, and tip fire-polished to resistances less than 4 MW. Series resistance of 3–9 MΩ was estimated from the whole-cell parameters circuit. Series resistance compensation (of < 90%) was used as needed to constrain voltage error to less than 10 mV, lag was 10 *μ*s. Cell capacitances were 4–15 pF. Capacitance and Ohmic leak were subtracted using a P/5 protocol. Output was low-pass filtered at 10 kHz using the amplifier’s built-in Bessel and digitized at 100 kHz. Traces were filtered at 2 kHz for presentation. Intersweep interval was 2 s. HEK293 cells with less than 65 pA/pF current at +85 mV were excluded to minimize impact of endogenous K^+^ currents (53).). The average current in the final 100 ms at holding potential prior to the voltage step was used to zero-subtract each recording. Mean outward current (*I*_avg,step_) was amplitude between 90-100 ms post depolarization. Mean tail current was the current amplitude between 0.2-1.2 ms into the 0 mV step. 100 *μ*L of 100 nM GxTX-594 was flowed over cells with membrane resistance greater than 1 GΩ, pulses to 0 mV gauged the time course of binding, and the *G–V* protocol was run. Data with predicted voltage error, *V*_error_ ≥ 10 mV was excluded from analysis. *V*_error_ was tabulated using estimated series resistance post compensation (*R*_s,post_)

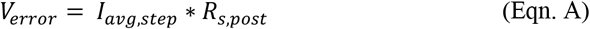

For *G–V* profiles cell membrane voltage (*V*_membrane_) was adjusted by *V*_error_ and *LJP*.

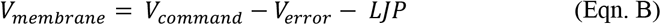

Tail currents were normalized by the mean current from 50 to 80 mV. Fitting was carried out using Igor Pro software, version 7 or 8 (Wavemetrics, Lake Oswego, OR) that employs nonlinear least squares curve fitting via the Levenberg-Marquardt algorithm. To represent the four independent and identical voltage sensors that must all activate for channels to open, *G–V* relations were individually fit with a 4^th^ power Boltzmann

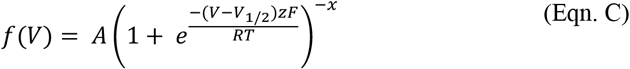

where *f*(V) is normalized conductance (*G*), *A* is maximum amplitude, *x* is the number of independent identical transitions required to reach full conductance (for a 4^th^ power function, *x*=4), *V*_1/2_ is activation midpoint, *z* is the valence in units of elementary charge (*e*_0_), *F* is the Faraday constant, *R* is the ideal gas constant, and *T* is absolute temperature. The half-maximal voltage (*V*_Mid_) for 4^th^ power functions is

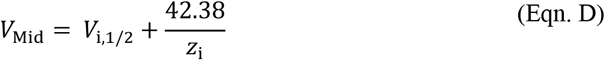

Reconstructed Boltzmann curves use average *z*_i_ and *V*_1/2_ ± SD. The minimum Gibbs free energy (Δ*G*_AMIGO1_) that AMIGO1 imparts to conductance, was tabulated as

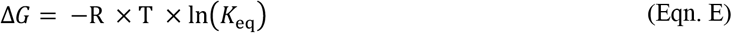

Here *R* = 0.00199 kcal/(K•mol) and *T* = 298K. *K*_eq_, or the equilibrium constant of channel opening, was approximated by 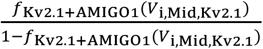 where *f*_Kv2.1+AMIGO1_(*V*_i,Mid,Kv2.1_) is the reconstructed relative conductance of Kv2.1 + AMIGO1 at *V*_i,Mid_ of Kv2.1–control cells (Table 1).

**Table 1.** Fourth order Boltzmann parameters for G–V relationships. Average *V*_i,1/2_, *V*_i,Mid_, and *z*_i_ values were derived from a 4^th^ order Boltzmann fits (Eqn. C) of *n* individual cells. All values are given ± SEM. Brown-Forsythe and Welch (appropriate for differing SD) ANOVA test with a Dunnett’s T3 multiple comparisons p-values: AB: 0.046. AC: 0.64. DE: 0.75. DF: 0.91. Unpaired, two-tailed t-test p-values: GH: 0.012. IJ: 0.95. KL: 0.0051. MN: 0.21. OP: 0.00018. QR: 0.19. Δ*G*_AMIGO1_ from Eqn. E, at *V*_i,Mid_ for Kv2.1 + GFP.

| | *G*–*V* fit parameters | | | | $\Delta G_{AMIGO1}$ (kcal/mol) |
| --- | --- | --- | --- | --- | --- |
| | $V_{i,1/2}$ (mV) | $V_{i,Mid}$ (mV) | $z_i$ ($e_0$) | $n$ | (Eqn. E) |
| **HEK293 cells** |  |  |  |  |  |
| mKv2.1 + GFP | -26.8 ± 3.0 | -1.7 ± 1.4 <sup>A</sup> | 1.79 ± 0.17 <sup>D</sup> | 7 | -0.31 |
| mKv2.1+ AMIGO1 + GFP | -30.9 ± 0.8 | -7.4 ± 1.8 <sup>B</sup> | 1.95 ± 0.16 <sup>E</sup> | 14 |  |
| mKv2.1 + SCNβ1 + GFP | -24.8 ± 1.5 | 0.2 ± 1.8 <sup>C</sup> | 1.720 ± 0.074 <sup>F</sup> | 8 |  |
| **Kv2.1–CHO cells** |  |  |  |  |  |
| rKv2.1 + GFP | -33.4 ± 1.7 | -1.8 ± 1.2 <sup>G</sup> | 1.411 ± 0.070 <sup>I</sup> | 20 | -0.28 |
| rKv2.1+ AMIGO1–YFP | -42.0 ± 3.3 | -7.6 ± 1.8 <sup>H</sup> | 1.40 ± 0.11 <sup>J</sup> | 19 |  |
| **Kv2.1–CHO cells + Mg<sup>2+</sup>** |  |  |  |  |  |
| rKv2.1 + GFP | -13.8 ± 1.8 | 17.6 ± 2.2 <sup>K</sup> | 1.51 ± 0.11 <sup>M</sup> | 18 | -0.37 |
| rKv2.1+ AMIGO1–YFP | -16.3 ± 1.5 | 10.2 ± 1.0 <sup>L</sup> | 1.682 ± 0.082 <sup>N</sup> | 23 |  |
| **Kv2.1–CHO cells + GxTX–594** |  |  |  |  |  |
| rKv2.1 + GFP | 26.8 ± 2.9 | 73.2 ± 3.8 <sup>O</sup> | 1.03 ± 0.11 <sup>Q</sup> | 13 | -0.77 |
| rKv2.1+ AMIGO1–YFP | 12.9 ± 4.4 | 50.9 ± 2.8 <sup>P</sup> | 1.27 ± 0.14 <sup>R</sup> | 12 |  |

Activation time constants (*τ*_act_) and sigmoidicity values (*σ*) (54) were derived by fitting 10-90% current rise with

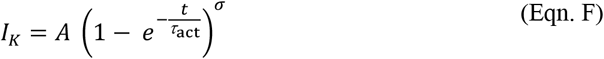

Where current at end of step, *I*_avg,step_, was set to 100%. *t* = 0 was adjusted to 100 *μ*s after voltage step start to correct for filter delay and cell charging. Deactivation time constants (*τ*_deact_) were from fitting 1 to 100 ms of current decay during 0 mV tail step with an exponential function

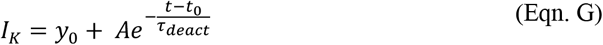

Reported *τ*_deact_ was the average after steps to +10 mV to +120 mV or +50 mV to +120 mV in GxTX–594. Kv2.1 deactivation kinetics became progressively slower after establishment of whole-cell mode, similar to Shaker deactivation after patch excision (55). Due to the increased variability of deactivation kinetics expected from this slowing phenomenon, deactivation kinetics were not analyzed further.

### On-cell single channel K^+^ currents

Single channel recordings were made from on-cell patches, to avoid Kv2.1 current rundown that occurs after patch excision (56). Methods same as whole-cell K^+^ ionic currents unless noted. While cells selected for recording had AMIGO1–YFP fluorescence apparent at the surface membrane, we cannot be certain each single Kv2.1 channel interacted with AMIGO1. Solutions: Kv2.1–CHO single channel internal (in mM) 155 NaCl, 50 HEPES, 20 KOH, 2 CaCl_2_, 2 MgCl_2_, 0.1 EDTA, adjusted to pH 7.3 with HCl, 347 mOsm. Kv2.1–CHO single channel external (in mM) 135 KCl, 50 HEPES, 20 KOH, 20 NaOH, 2 CaCl_2_, 2 MgCl_2_, 0.1 EDTA, adjusted to pH 7.3 with HCl, 346 mOsm: LJP −3.3 mV with Kv2.1–CHO single channel internal. Thick-wall borosilicate glass (BF150-86-7.5HP; Sutter Instruments) was pulled, Sylgard-coated and fire–polished, to resistances >10 MΩ. Analysis methods were same as prior (5) unless noted. To subtract capacitive transients, traces without openings were averaged and subtracted from each trace with single-channel openings. Peaks in single channel amplitude histograms were fit to half maximum with a Gaussian function to define single channel opening level for idealization by half-amplitude threshold. Open dwell times were well described by a single exponential component which was used to derived *τ*_closing_. Average open dwell times were also described as the geometric mean of all open dwell times. Closed dwell times appeared to have multiple exponential components and were solely described as the geometric mean of all closed dwell times.

### Whole–cell gating current measurements

Methods same as whole-cell K^+^ ionic currents unless noted. Solutions: gating current internal (in mM) 90 NMDG, 1 NMDG-Cl, 50 HEPES, 5 EGTA, 50 NMDG-F, 0.01 CsCl, adjusted to pH 7.4 with methanesulfonic acid, 303 mOsm. Gating current external (in mM) 150 TEA-Cl, 41 HEPES, 1 MgCl_2_ · 6 H_2_O, 1.5 CaCl_2_, adjusted to pH to 7.3 with NMDG, 311 mOsm: LJP −3.3 mV with gating current internal. To avoid KCl contamination of the recording solution from the pH electrode, pH was determined in small aliquots that were discarded. Cells were resuspended in Kv2.1–CHO external and washed in the recording chamber with 10 mL gating current external. Pipettes has resistances of 6-14 MΩ. Series resistances were 14-30 MΩ and compensated 50%. Cell capacitances were 6-10 pF. *V_error_* was negligible (< 1 mV). P/5.9 leak pulses from −133 mV leak holding potential. An early component ON gating charge movement was quantified by integrating ON gating currents in a 3.5 ms window (*Q*_ON,fast_) following the end of fast capacitive artifacts created from the test voltage step (which usually concluded 0.1 ms following the voltage step). The slow tail of the ON charge movement is difficult to accurately integrate in these cells, making the cutoff point arbitrary. This 3.5 ms integration window resulted in a more positive *Q*_ON,fast_–*V* midpoint than with a 10 ms window (5), and more positive midpoint than the *G–V* relation. Differences in gating current solutions compared to prior studies may also contribute to the different midpoints reported (4, 5, 57). Currents were baseline-subtracted from 4 to 5 ms into step. *Q*_OFF_ was determined by integration of OFF charge movement in a 9.95 ms window after capacitive artifacts (usually 0.1 ms). Currents were baseline-subtracted from 10 to 20 ms into the step. Gating charge density *fC/pF* was normalized by cell capacitance. *Q–V* curves normalized to average from 100-120 mV. *Q–V* relations were individually fit with a 1^st^ power Boltzmann (Eqn. C., *x*=1). Time constants (*τ*_ON_) were determined from a double-exponential fit function

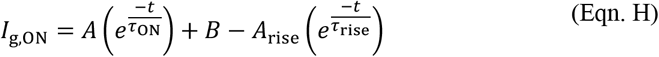

*τ*_rise_ was not used in analyses. *I*_g,OFF_ was not well fit by Eqn. H and *τ*_OFF_ was not analyzed. The voltage-dependence of the forward voltage sensor activation (*α*) rate was determined by fitting the average *τ*_ON_–*V* weighted by the standard error

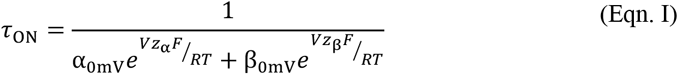

Reverse rates were not analyzed. Energy of AMIGO1 impact on the activation rate of all 4 voltage sensors (Δ*G^‡^AMIGO1*) was

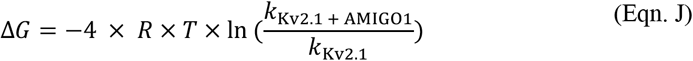

where *k* = *α*_0mv_. Estimates of Δ*G*_AMIGO1_ from *Q–V* relations were with Eqn. E or

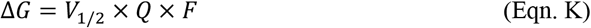

Here *F* = 23.06 kcal/V • mol • *e*_0_. *Q* was either *z_g_* from fits or 12.5 *e*_0_ as determined from a limiting slope analysis of the Kv2.1 open probability-voltage relation (3). *V*_1/2_ was either *V*_g,Mid_ or a median voltage (*V*_g,Med_) as calculated from integration above and below *Q*_OFF_–*V* relations using a trapezoidal rule (58).

### Fluorescence imaging

Images were obtained with an inverted confocal/airy disk imaging system with a diffraction grating separating 400-700 nm emission into 9.6 nm bins (Zeiss LSM 880, 410900-247-075) run by ZEN black v2.1. Laser lines were 3.2 mW 488 nm, 1.2 mW 514 nm, 0.36 mW 543 nm, 0.60 mW 594 nm. Images were acquired with a 1.4 NA 63x (Zeiss 420782-9900-799), 1.3 NA 40x (Zeiss 420462-9900-000), or 1.15 NA 63x objectives (Zeiss 421887-9970-000). Images were taken in either confocal or airy disk imaging mode. The imaging solution was Kv2.1–CHO external supplemented with 0.1% bovine serum albumin and 10 mM glucose. Temperature inside the microscope housing was 24-28 °C. Representative images had brightness and contrast adjusted linearly.

#### Concentration-effect imaging

Cells plated on coverslips were washed 3x with imaging solution then mounted on an imaging chamber (Warner Instruments, RC-24E) with vacuum grease. 100 *μ*L GxTX–594 dilutions were applied for 10 minutes, then washed-out by flushing 10 mL at a flow rate of ~1 mL / 10 sec. 15 minutes after wash–out, the next GxTX–594 concentration was added. Airy disk imaging, 1.4 NA 63x objective (Zeiss 420782-9900-799), 0.13 *μ*m pixels, 0.85 *μ*s dwell, 5 sec frame rate. YFP excitation 488 nm 2% power, emission 495-550 nm. GxTX–594 excitation 594 nm 2% power, emission 495-620 nm. Intensities extracted using FIJI (59). ROIs drawn around groups of cells ± YFP fluorescence. Dissociation constant (*K_d_*) fit with fluorescence intensity at 0 nM GxTX–594 set to 0 with

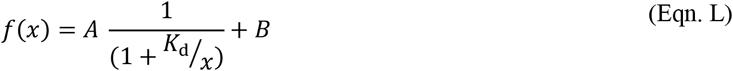

*Voltage clamp fluorimetry* was conducted as described (45). Briefly, 100 *μ*L 100 nM GxTX–594 in imaging external was applied for 10 minutes then diluted with 1 mL Kv2.1–CHO external for imaging. Airy disk imaging, 1.15 NA 63x objective (Zeiss 421887-9970-000), 0.11 *μ*m pixels, 0.85 *μ*s dwell, 2x averaging, 1 sec frame rate. GxTX–594 excitation 594 nm 1% power, emission 605nm long-pass. Cells with obvious GxTX–594 labeling were whole-cell voltage-clamped. Voltage clamp fluorimetry internal (in mM) 70 mM CsCl, 50 mM CsF, 35mM NaCl, 1 mM EGTA, 10 mM HEPES, adjusted to pH 7.4 with CsOH, 310 mOsm: LJP −5.3 mV with Kv2.1–CHO external. Pipettes from thin-wall glass were less than 3.0 MΩ. Cells were held at −100 mV for 30 images and stepped to +35 mV until fluorescence change appeared complete. Intensity data was extracted using Zen Blue from ROIs drawn around apparent surface membrane excluding pipette region. For presentation, fluorescence intensity traces were normalized from minimum to maximum. Rate of GxTX–594 dissociation (*k_ΔF_*) was fit with a monoexponential function (Eqn. G), and *K_eq_* for resting vs. activated voltage sensors was calculated as described (45). Δ*G*_AMIGO1_ from with Eqn. J where *k* = *K_eq_*.

*Environment-sensitive fluorescence imaging* with GxTX Ser13Pra(JP) and GxTX Lys27Pra(JP). Cells were incubated in 100 *μ*L of GxTX(JP) solution for 5-10 minutes then washed with imaging solution. Spectral confocal imaging, 1.4 NA 63x objective, 0.24 *μ*m pixels, 8.24 *μ*s dwell, 2x averaging. YFP excitation 514 nm. GxTX Ser13Pra(JP) excitation 594 nm. GxTX Lys27Pra(JP) excitation 543 nm. Fluorescence counts extracted in Zen Blue. JP emission spectra were fit with two-component split pseudo-Voigt functions (46) using the curve fitting software Fityk 1.3.1 (https://fityk.nieto.pl/), which employed a Levenberg-Marquardt algorithm. Goodness of fit was determined by root-mean-squared deviation (*R*^2^) values, which are listed in Supplemental Table 2 along with the parameters of each component function. To avoid YFP overlap, fittings for spectra from cells expressing AMIGO1–YFP include emission data points from 613-700 nm for GxTX Ser13Pra(JP) and 582-700 nm for GxTX Lys27Pra(JP). Fittings for JP spectra from cells without AMIGO1-YFP included all data from 550-700 nm.

### Experimental Design and Statistical Treatment

Independent replicates (*n*) are individual cells pooled over multiple transfections. The *n* from each transfection for each figure are listed in Supplemental Tables 3 and 4. In each figure panel, control and test cells were plated side by side from the same suspensions, transfected side by side, and the data was acquired from control and test cells in an interleaved fashion. Identity of transfected constructs was blinded during analysis. ANOVA analysis of transfection- or acquisition date-dependent variance of Boltzmann fit parameters and PCC/COV did not reveal a dependence, and all *n* values were pooled. Statistical tests were conducted with Prism 9 (GraphPad Software, San Diego, CA), details in figure legends.

## Results

### AMIGO1 shifts the midpoint for activation of Kv2.1 conductance

Voltage-clamp recordings from cotransfected HEK293 cells indicate that mouse AMIGO1 shifts the *G–V* relation of mouse Kv2.1 by −5.7 ± 2.3 mV (SEM) (Supplemental Fig. 1). This shift was similar to the −6.1 mV ± 1.6 mV shift reported of rat Kv2.1–GFP by human AMIGO1–mRuby2 (23), and smaller than the −15.3 mV (no error listed) shift of mouse Kv2.1– GFP by mouse AMIGO1 (22). This small effect of AMIGO1 was similar to the cell-to-cell variability in our recordings. We suspected that endogenous voltage-activated conductances of HEK293 cells (53, 60) and variability inherent to transient co-transfection could increase variability. To minimize possible sources of cell-to-cell variability, further experiments were with a Chinese Hamster Ovary K1 cell line with inducible rat Kv2.1 expression (Kv2.1–CHO) transfected with a YFP-tagged mouse AMIGO1. Inducible Kv2.1 expression permits tighter control of current density (49) and fluorescence tagging of AMIGO1 permits visualization of protein expression and localization. Unlike HEK293 cells, CHO cells lack endogenous voltage-gated K^+^ currents (61).

As expression systems can influence auxiliary protein interactions with ion channels (62–66), we assessed Kv2.1–AMIGO1 association in these CHO cells. We evaluated two hallmarks of Kv2.1 and AMIGO1 association: Kv2.1 reorganization of AMIGO1, and AMIGO1 / Kv2.1 colocalization (22, 23, 28).

In HEK293 cells, heterologously expressed AMIGO1 localization is intracellular and diffuse (23, 28). However, when co-expressed with Kv2.1, AMIGO1 reorganizes into puncta with Kv2.1, similar to the expression patterns in central neurons (23, 28). To determine whether Kv2.1 reorganizes AMIGO1 in Kv2.1–CHO cells, the degree of AMIGO1–YFP reorganization was quantified using the Coefficient of Variation (COV), which captures non–uniformity of YFP localization (67). COV was quantified following the limited 1.5 h Kv2.1 induction period used in whole-cell and single channel K^+^ current recordings and the prolonged 48 h induction period used for gating current recordings or imaging studies. COVs were compared against an uninduced control (0 h induction) and against an engineered protein, ChroME-mRuby2, which contains the Kv2.1 PRC trafficking sequence, but lacks the Kv2.1 voltage sensing and pore forming domains (51, 52). COVs were evaluated from the glass-adhered, basal membrane where evidence of reorganization is most notable (Fig. 1). Both COV_1.5h_ and COV_48h_ were greater than the COV_0h_ or COV_ChroME-mRuby_ control. This result is consistent with Kv2.1 and AMIGO1 association in CHO cells.

**Figure 1.**
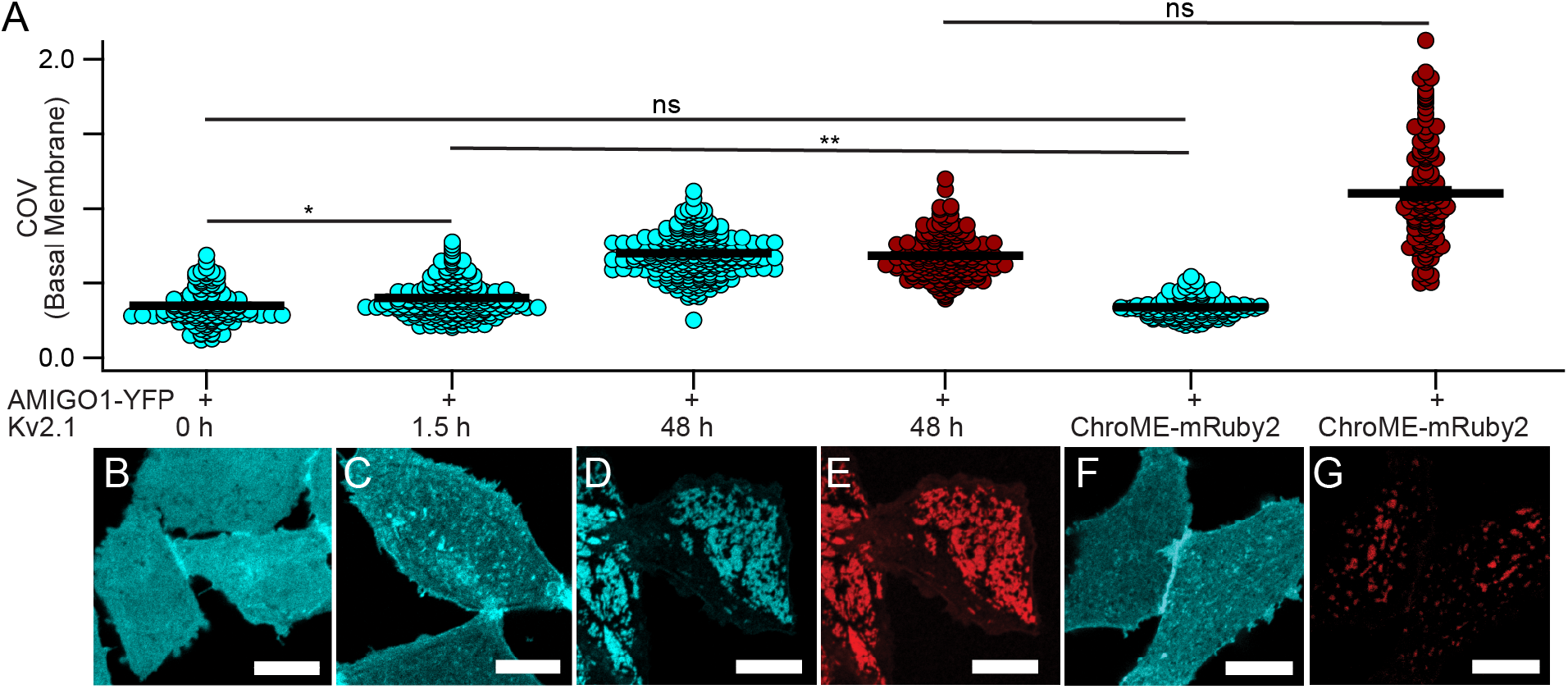
Kv2.1 reorganizes AMIGO1 in CHO cells. (**A**) Coefficient of variation of fluorescence from AMIGO1–YFP (blue circles), GxTX–594 (red circles), or ChroME-mRuby2 (red circles). Bars are mean ± SEM. COV measurements were calculated from confocal images acquired from the glass–adhered basal membrane of the cell (exemplar confocal images in B-G). All cells were transfected with AMIGO1–YFP 48 h prior to imaging. COV from individual cells (n) were pooled from 4 separate transfections for each experimental condition. AMIGO1–YFP fluorescence from cells (**B**) not induced for Kv2.1 expression (COV_0h_ = 0.3492 ± 0.0098, n = 134), (**C**) induced 1.5 h (COV_1.5h_ = 0.4013 ± 0.0077, n = 217), (**D**) induced 48 h (COV_48h_ = 0.6984 ± 0.0083, n = 277). (**E**) GxTX–594 labeling from panel D (COV_48h(GxTX–594)_ = 0.6822 ± 0.010, n = 197). (**F**) AMIGO1–YFP fluorescence from CHO cells which lack Kv2.1 co-transfected with ChroME-mRuby2 (COV_lack_ = 0.3377 ± 0.0059, n = 125). (**G**) ChroME-mRuby2 fluorescence from panel F (COV_(ChroME-mRuby2)_ = 1.102 ± 0.030, n = 128). Scale bars 10 μm. (**Statistics**) Outliers removed using ROUT, Q = 1%. Ordinary one-way ANOVA with multiple comparisons. P-values: COV_0h_COV_1.5h_: p = 0.0467; COV_0h_COV_lack_: p = 0.9936; COV_1.5h_COV_lack_: p = 0.0081; COV_48h(GxTX–594)_ COV_(ChroME-mRuby2)_: p =0.9010. All other p-values ≤ 0.0001.

As an additional measure of whether Kv2.1 reorganizes AMIGO1 in Kv2.1–CHO cells, we assessed AMIGO1–YFP and Kv2.1 colocalization using the Pearson’s correlation coefficient (PCC) (68). Surface-expressing Kv2.1 on live cells was labeled with GxTX Ser13Cys(Alexa594), a conjugate of a voltage sensor toxin guangxitoxin-1E derivative with a fluorophore, abbreviated as GxTX–594 (45). As auxiliary subunits can impede binding of toxins to voltage-gated ion channels (69), we tested whether AMIGO1 impacted GxTX–594 binding to Kv2.1. Under conditions where AMIGO1 modulates most, if not all, Kv2.1 voltage sensor movements (Fig. 6, 7), we found no evidence that AMIGO1 impedes GxTX–594 binding to Kv2.1 (Supplemental Fig. 5). Colocalization between AMIGO1–YFP and GxTX–594 was apparent as PCC_48h_, measured from the glass-adhered basal membrane, was greater than the negative control, PCC_ChroME-mRuby2_ (Fig. 2B). With a limited 1.5 h induction, GxTX–594 was difficult to detect at the glass-adhered membrane, so we moved the confocal imaging plane further from the cover glass to image Kv2.1 on apical cell surfaces where GxTX–594 labeling was more apparent. On these apical surfaces, PCC_1.5h_ and PCC_48h_ were greater than PCC_0h_ (Fig. 2A), consistent with some colocalization of AMIGO1–YFP and Kv2.1. The weakly significant increase of the PCC_1.5h_ compared to PCC_0h_ is consistent with some colocalization. Disproportionate expression can skew PCC values (70), and the limited GxTX–594 signal is expected to depress the PCC_1.5h_ value. Similarly, the lower PCC_48h_ values were associated with either minimal or exceptionally bright AMIGO1–YFP signal. Overall, we see no sign of Kv2.1 channels lacking colocalized AMIGO1 in cells with high levels of AMIGO1 expression. Altogether, the reorganization and colocalization indicate that AMIGO1–YFP and Kv2.1 interact in the CHO cells used for K^+^ current recordings and for gating current measurements.

**Figure 2.**
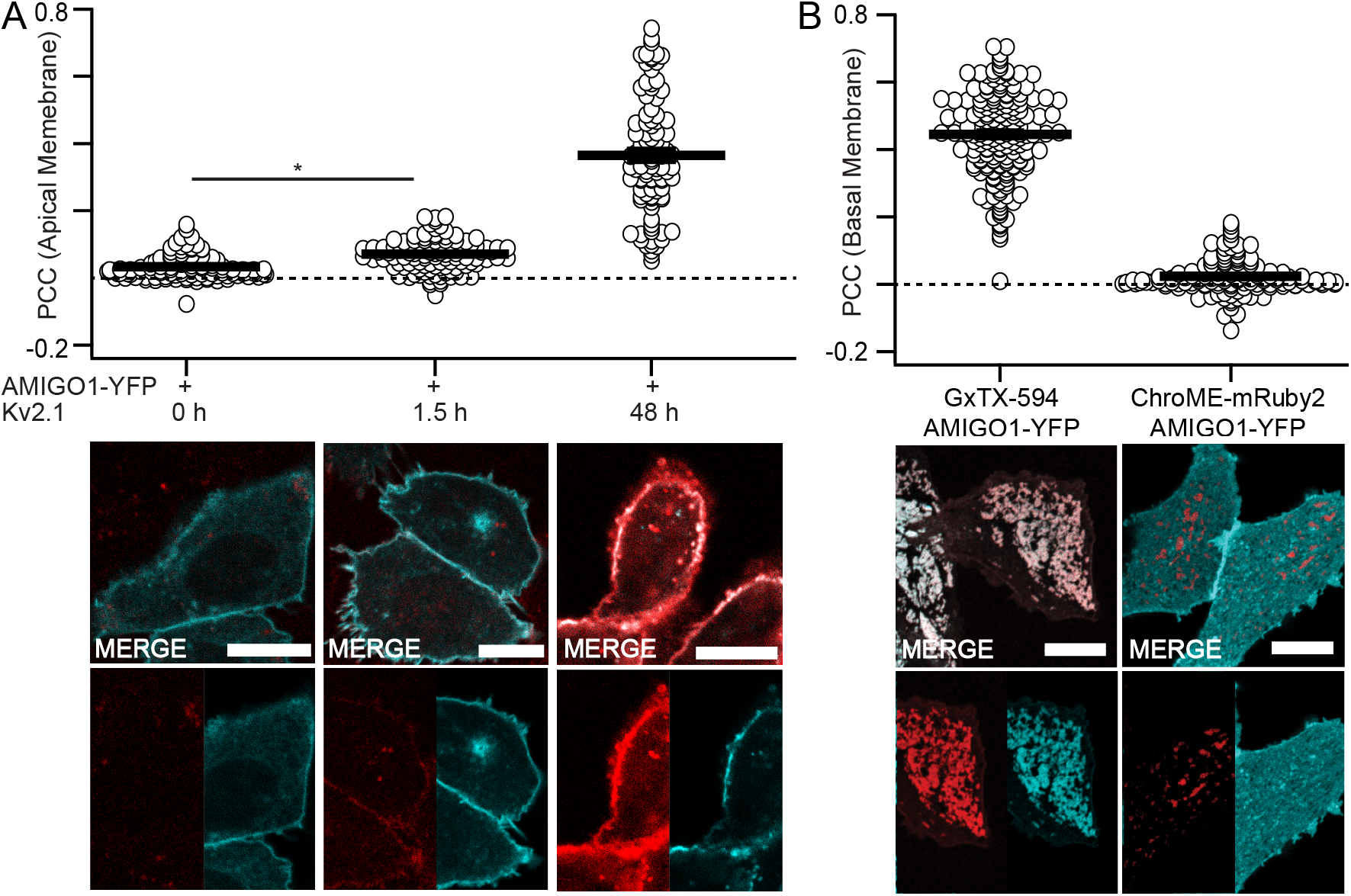
AMIGO1 colocalizes with Kv2.1 in CHO cells. (**A**) Costes thresholded, Pearson’s colocalization between AMIGO1–YFP and GxTX–594 at cell membrane following, from left to right, 0, 1.5, or 48 h of Kv2.1 induction (exemplar confocal images in B-J below). Mean ± SEM (one-tailed ≥ 0 t-test): PCC_0h_ = 0.0321 ± 0.0033, (p < 0.0001), *n* = 101; PCC_1.5h_ = 0.0718 ± 0.0042, (p < 0.0001), *n* = 118; and PCC_48h_ = 0.365 ± 0.017, (p < 0.0001), *n* = 101. (**B**) Costes thresholded, Pearson’s colocalization between (left to right) AMIGO1–YFP/GxTX–594 and AMIGO1–YFP/ChroME-mRuby2 at the glass-adhered basal membrane of the cell. Exemplar images are the same as in Fig. 1 D-G. From left to right: PCC_GxTX–594_ = 0.4449 ± 0.0090, (p < 0.0001), *n* = 195; PCC_ChroME-mRuby2_ = 0.0242 ± 0.0045, (p < 0.0001), *n* = 129. Image panels with merge overlays (white) of GxTX–594 (red) and AMIGO1–YFP (cyan) correspond to conditions above. All scale bars are 10 μm. (**Statistics**) Outliers were removed using ROUT, Q = 1%. Ordinary one-way ANOVA with multiple comparisons. P-values: PCC_0h_PCC_1.5h_: p = 0.346; PCC_1.5h_PCC_ChroME-mRuby2_: p = 0.0025; PCC_0h5h_PCC_ChroME-mRuby2_: p = 0.9777. All other p-values were ≤ 0.0001.

### AMIGO1 shifts the midpoint of activation of Kv2.1 conductance in CHO cells

To determine whether AMIGO1 affected the macroscopic K^+^ conductance in Kv2.1– CHO cells, we conducted whole-cell voltage clamp recordings. Cells were transfected with GFP (Kv2.1–control cells) or with AMIGO1–YFP (Kv2.1 + AMIGO1 cells) and identified for whole-cell voltage clamp based on the presence of cytoplasmic GFP fluorescence or plasma membrane-associated YFP fluorescence, respectively (Fig. 3A). Macroscopic ionic current recordings were made in whole-cell voltage-clamp mode and K^+^ conductance was measured from tail currents (Fig. 3B, C). In expectation of small AMIGO1 effects relative to cell-to-cell variation, recordings from control cells and AMIGO1 cells were interleaved during each day of experiments and cell identity was blinded during analysis. *G–V* relations were fit with a 4^th^ power Boltzmann function (Eqn. C) (Fig. 3D, E, F) and average midpoints of half-maximal conduction (*V*_i,Mid_) and steepness equivalents (*z_i_*) were determined (Table 1). In Kv2.1–control cells, the average *V*_i,Mid_ was −1.8 mV (Fig. 3H), consistent with prior reports of *V*_i,Mid_ ranging from −3 mV to +8 mV in CHO cells (4, 23, 47, 71). Cell-to-cell variation in *V*_i,Mid_ remained notable between Kv2.1–CHO cells, with variation in *V*_i,Mid_ on par with other reports (see *Discussion/Limitations*). The range of *V*_i,Mid_ values of Kv2.1 + AMIGO1 cells overlapped with Kv2.1–control cells (Fig. 3H), yet the average *V*_i,Mid_ was negatively shifted by −5.7 ± 2.2 mV (SEM), similar to Δ*V*_i,Mid_ from mouse Kv2.1 in HEK293 cells (Table 1). No effect on *z*_i_ was observed. We also tested AMIGO2 and AMIGO3 on Kv2.1, and found they colocalize and induce Δ*V*_i,Mid_ shifts similar to those reported from HEK293 cells by Maverick and colleagues (23) (Supplemental Fig. 3, 4), indicating that the small *G–V* shifts by the AMIGO proteins are robust across different experimental preparations.

**Figure 3.**
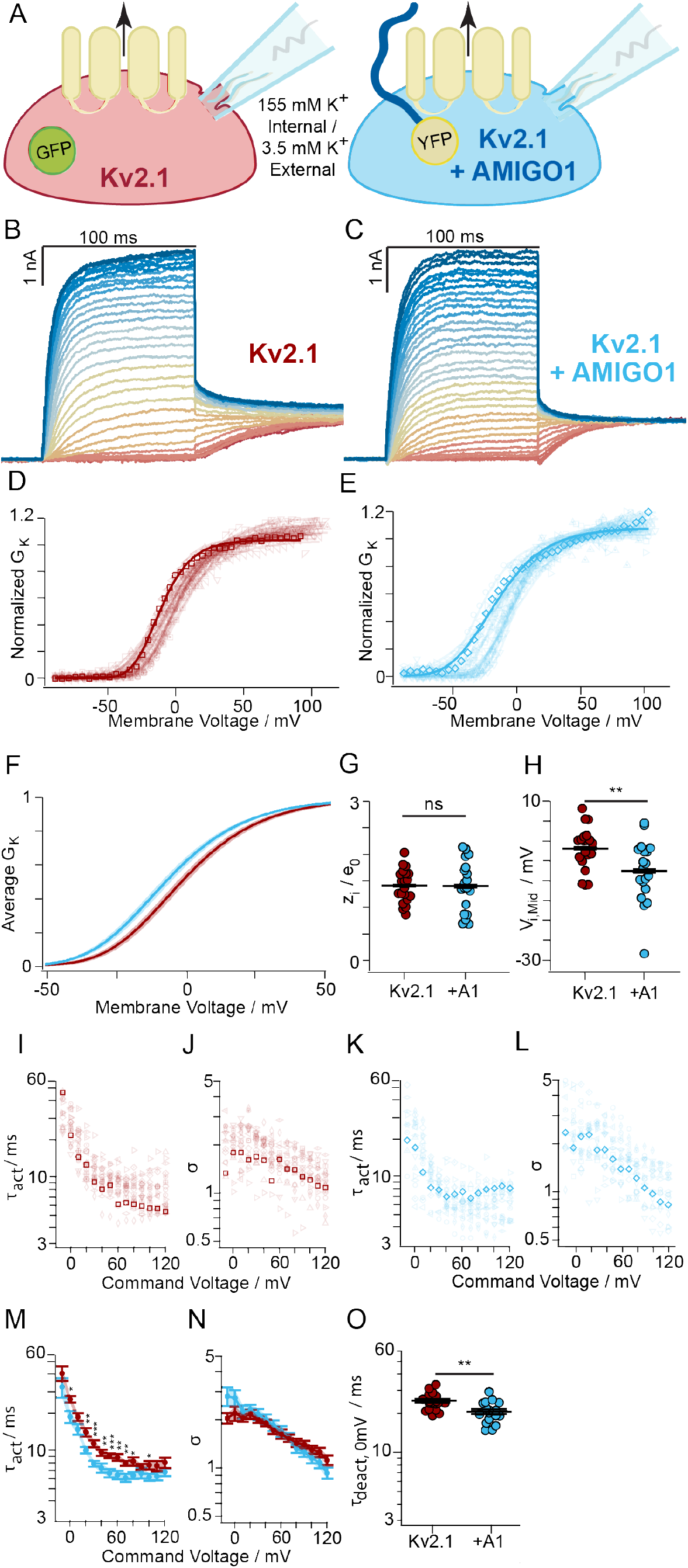
AMIGO1 shifts the midpoint and speeds activation of the Kv2.1 conductance in CHO cells. (**A**) Experimental set up: Whole-cell K^+^ currents (arrow) from Kv2.1–CHO transfected with GFP (red) or AMIGO1–YFP (blue). (**B**, **C**) Representative Kv2.1–control (6.0 pF) or Kv2.1 + AMIGO1 (14.5 pF) cell. 100 ms voltage steps ranging from −80 mV (dark red trace) to +120 mV (dark blue trace) in 5 mV increments and then to 0 mV for tail currents. Holding potential was −100 mV. Data points from representative cells are bolded in analysis panels. (**D**, **E**) Normalized tail *G–V* relationships for Kv2.1–control or Kv2.1 + AMIGO1 cells. Symbols correspond to individual cells. Lines are 4^th^ order Boltzmann fits (Eqn. C). (**F**) Reconstructed Boltzmann fits from average Vi,Mid and zi (Table 1). Shading Vi,Mid ± SEM. (**G**) Steepness and (**H**) midpoint of fits. (**I**, **K**) *τ*_act_ and (**J**, **L**) σ from fits of Eqn. F to activation (**M**) Mean *τ*_act_ and (**N**) σ. (**O**) *τ*_deact_ fits of Eqn. G to 0 mV tails: Kv2.1–control 24.9 ± 3.6 ms, Kv2.1+AMIGO1 20.6 ± 3.8 ms. Unpaired t-test p > 0.5 between 0 mV *τ*_act_ and *τ*_deact_ for Kv2.1–control and Kv2.1 + AMIGO1. All other statistics in Table 1. ***: p = ≤0.001, **: p = ≤0.01, *: p = ≤0.05, ns: not significant. Bars are mean ± SEM.

To test if AMIGO1 also alters the rate of activation of Kv2.1 conductance, we analyzed activation kinetics. The 10-90% of the rise of Kv2.1 currents following a voltage step (Fig. 3A, B) was fit with the power of an exponential function (Eqn. F) for sigmoidicity (*σ*) which quantifies delay before current rise, and activation time constant (*τ*_act_). *σ* was not significantly altered by AMIGO1 (Fig. 3J, L, N), suggesting that the Kv2.1 activation pathway retains a similar structure with AMIGO1 (5). At a subset of voltages less than +70 mV, AMIGO1 expression accelerated activation, decreasing *τ*_act_ (Fig. 3I, K, M), consistent with results of Maverick and colleagues (23). Following the +10 to +120 mV activating steps, time constants of tail current decay at 0 mV were similar to *τ*_act_ at 0 mV (Fig. 3O, Eqn. G). A prior study found no impact of AMIGO1 on Kv2.1 deactivation kinetics at −40 mV (23), and deactivation is not studied further here. A model of Kv2.1 activation kinetics suggests that voltage sensor dynamics influence *τ*_act_ below ~+70 mV, and that at more positive voltages a slow pore opening step limits kinetics (5). This analysis suggests that AMIGO1 accelerates activation kinetics only in the voltage range which is sensitive to voltage sensor dynamics.

### Effects of AMIGO1 on pore opening conformational changes were not apparent in single channel recordings

To more directly assess whether the pore opening step of the Kv2.1 activation pathway is modulated by AMIGO1, we analyzed pore openings of single Kv2.1 channels during 1 s long recordings to 0 mV (Fig. 4A, B). At 0 mV we expect >85% of all Kv2.1–control voltage sensors or >95% of all Kv2.1–AMIGO1 voltage sensors (Fig. 6T) to activate in less than 2 ms (Fig. 6N), such that the majority of single channel openings represent stochastic fluctuations between a closed and open conformation of the pore. Neither the single channel current amplitude (Fig. 4C, D, E) nor the intra–sweep open probability (Fig. 4F) were significantly impacted by AMIGO1. AMIGO1 did not significantly impact the single channel open or closed dwell times (Fig. 4G-L). These results constrain any impact of AMIGO1 on Kv2.1 pore opening to be smaller than the variability in these single channel measurements.

**Figure 4.**
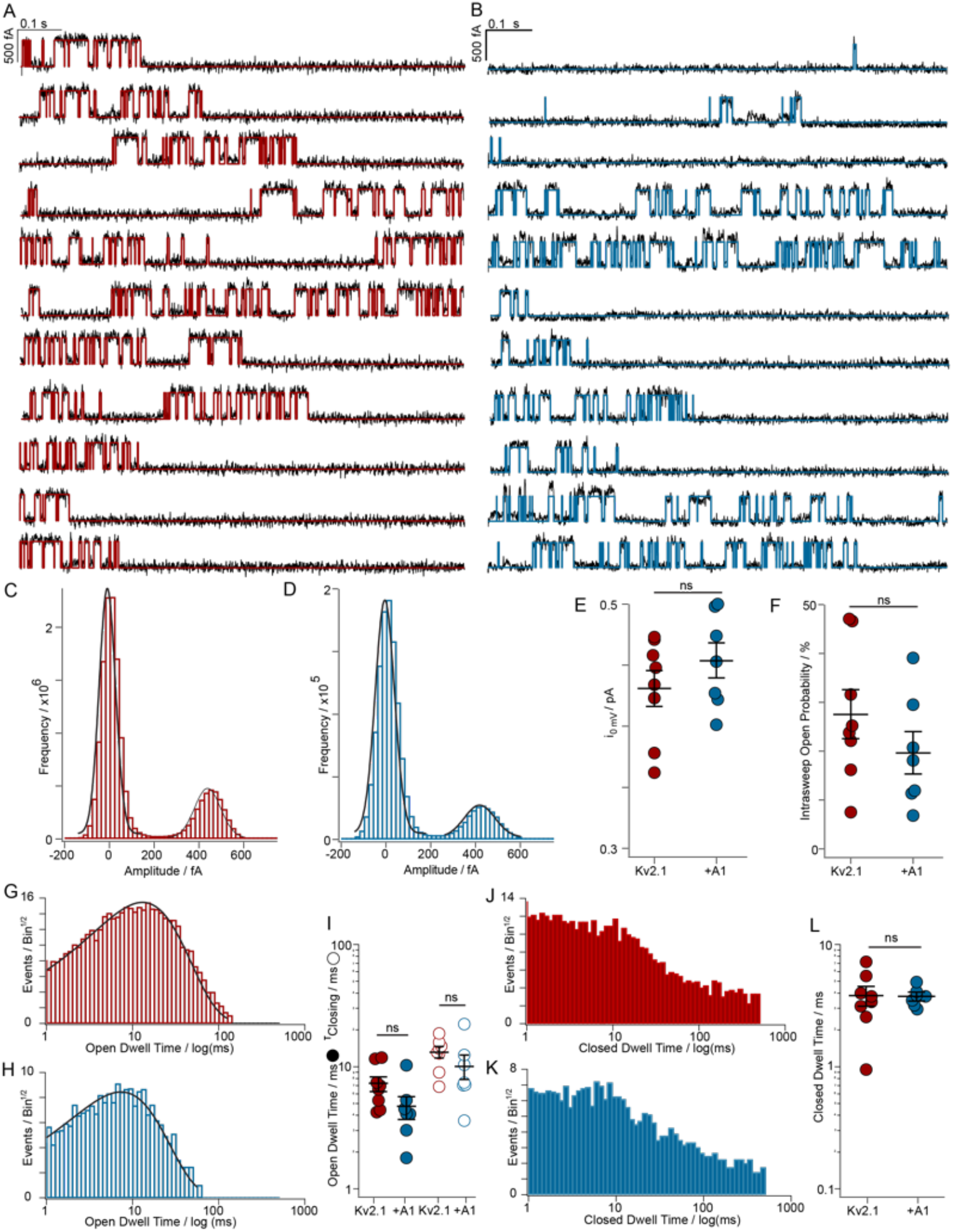
Effects of AMIGO1 on pore opening conformational changes were not apparent in single channel recordings. (**A**) Representative single channel currents at 0 mV from Kv2.1–control and (**B**) Kv2.1 + AMIGO1. Red or blue lines are idealizations. (**C,D**) Amplitude histograms at 0 mV from the patches in A,B fit with Gaussians. (**E**) Mean single channel current amplitude: Kv2.1–control 0.43 ± 0.01 pA, Kv2.1 + AMIGO1 0.45 ± 0.02 pA. (**F**) Open probability from amplitude histograms: Kv2.1–control 28 ± 4.9%, Kv2.1 + AMIGO1 20 ± 4.2%. (**G**) Open dwell-time distributions and single exponential fits for a Kv2.1–control or (**H**) Kv2.1 + AMIGO1 patch. (**I**) Open dwell times from mean (filled circles) or exponential fit (hollow circles). Kv2.1–control: 13.0 ± 1.3 μs. Kv2.1 + AMIGO1: 9.98 ± 2.3 μs. (**J**) Closed dwell-time distributions and single exponential fit for a Kv2.1–control or (**K**) Kv2.1 + AMIGO1 patch. (**L**) Closed dwell times from mean.. Kv2.1–control: 3.80 ± 0.67 μs. Kv2.1 + AMIGO1: 3.73 ± 0.250 μs. ns = two-tailed t-test p-value > 0.05. Means ± SEM.

### A voltage sensor toxin enhances modulation of AMIGO1 on the Kv2.1 conductance

To test whether AMIGO1 modulation is dependent on voltage sensor dynamics, we altered voltage sensor movement with a voltage sensor toxin. GxTX binds to the voltage sensing domain of Kv2.1 (72), such that exit from the earliest resting conformation limits opening to more positive voltages (5). If AMIGO1 modulates voltage sensors, then GxTX might be expected to amplify the AMIGO1 effect. Alternately, if AMIGO1 acts directly on pore opening, the AMIGO1 impact on the pore opening equilibrium should persist, regardless of voltage sensor modulation. To distinguish between these possibilities, we measured AMIGO1 modulation in the presence of the imaging probe GxTX–594, which modulates Kv2.1 by the same mechanism as GxTX (45) and has a similar affinity for the resting conformation of Kv2.1 with or without AMIGO1 (Supplemental Fig. 5). We applied 100 nM GxTX–594 to cells and activated the Kv2.1 conductance. We note that the 100-ms activating pulses are much shorter than the >2 second time constants of GxTX–594 dissociation at extreme positive voltages (45) and during these short activating pulses we saw no evidence of GxTX–594 dissociation. The AMIGO1 Δ*V*_i,Mid_ of −22.1 ± 4.8 (SEM) with GxTX–594 was distinct from the AMIGO1 Δ*V*_i,Mid_ of −5.7 ± 2.2 mV (SEM) without GxTX–594 (p = 0.00018, unpaired, two-tailed t-test), indicating that GxTX–594 amplifies the impact of AMIGO1 on Kv2.1 conductance. We did not observe a significant effect of AMIGO1 on *τ*_act_ or *σ* in GxTX–594 (Fig. 5J-N). We calculated the impact of AMIGO1 on a pore opening equilibrium constant (*K*_eq_) at the midpoint of the Kv2.1 *G–V* relation and found a 3.7-fold bias towards a conducting conformation in 100 nM GxTX–594 versus a 1.4-fold bias under control conditions (Δ*G*_AMIGO1_ = −0.77 versus −0.28 kcal/mol respectively, Table 1). This result indicates that the impact AMIGO1 has on the Kv2.1 conductance is dependent on the dynamics of the activation path. Further, this result indicates that AMIGO1 opposes the action of GxTX–594, which stabilizes the earliest resting conformations of Kv2.1 voltage sensor. We also note that the more dramatic modulation by AMIGO1 with GxTX–594 verifies that most Kv2.1 channels are modulated by AMIGO1 in this cell preparation in which only a small impact on *V*_i,Mid_ was observed without GxTX–594 (Fig. 3).

**Figure 5.**
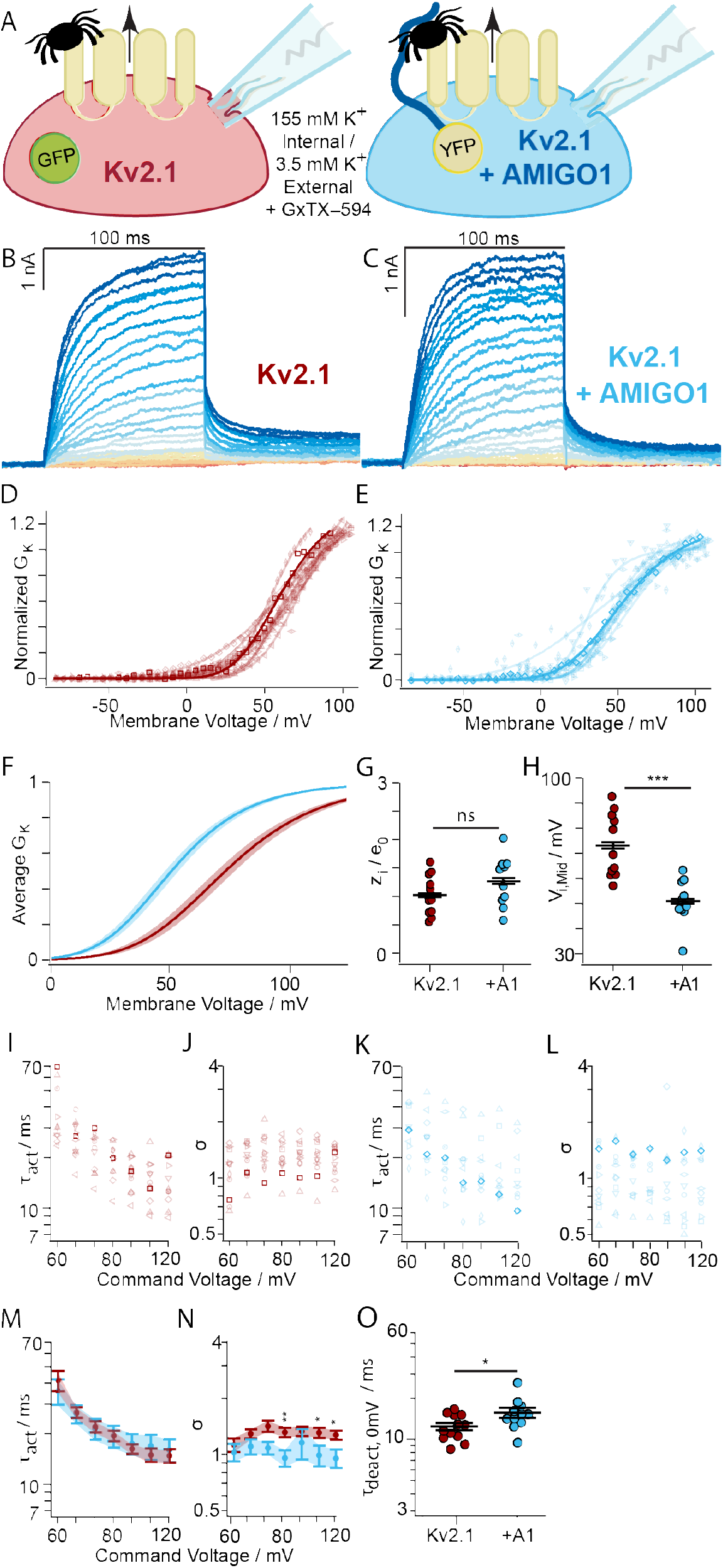
The voltage sensor toxin GxTX–594 enhances AMIGO1 modulation of Kv2.1 conductance. (**A**) Experimental set up: Whole-cell K^+^ currents (arrow) from Kv2.1–CHO transfected with GFP (red) or AMIGO1–YFP (blue). Cells were treated with 100 nM GxTX–594 (tarantulas). (**B, C**) Representative Kv2.1–control (6.0 pF) or Kv2.1 + AMIGO1 (14.5 pF) cell. Same voltage protocol and representations as Fig. 3. (**D, E**) Normalized *G–V* relationships (**F**) Reconstructed 4^th^ order Boltzmann fits from *V*_i,Mid_ and *z*_i_ in Table 1. Shading *V*_i,Mid_ ± SEM. (**G**) Steepness and (**H**) midpoint of fits. (**I**, **K**) *τ*_act_ and (**J**, **L**) *σ* from fits of Eqn. F to activation (**M**) Mean *τ*_act_ and (**N**) *σ*. (**O**) *τ*_deact_ fits of Eqn. G to 0 mV tails: Kv2.1 with GxTX–594 = 12.4 ± 2.7 ms. Kv2.1+AMIGO1 with GxTX–594 = 15.7 ± 4.2 ms. All other statistics in Table 1. ***: p ≤0.001, **: p = ≤0.01, *: p = ≤0.05, ns: not significant. Bars are mean ± SEM.

### AMIGO1 facilitates the activation of Kv2.1 voltage sensors

To determine if AMIGO1 affects voltage sensor movement, we measured gating currents (*I*_g_), which correspond to movement of Kv2.1 voltage sensors across the transmembrane electric field. Kv2.1–CHO cells were patch clamped in whole–cell mode in the absence of K^+^ (Fig. 6A) and given voltage steps to elicit gating currents (Fig. 6B, C). The resolvable ON gating currents (*I*_g,ON_) represent an early component of gating charge movement, but not all of the total gating charge; the later charge movements, which include any charge associated with the pore opening, move too slowly for us to resolve accurately in ON measurements (4, 5). If AMIGO1 acts solely through the pore we would not expect to detect an impact on early components of ON gating currents which occur before pore opening.

**Figure 6.**
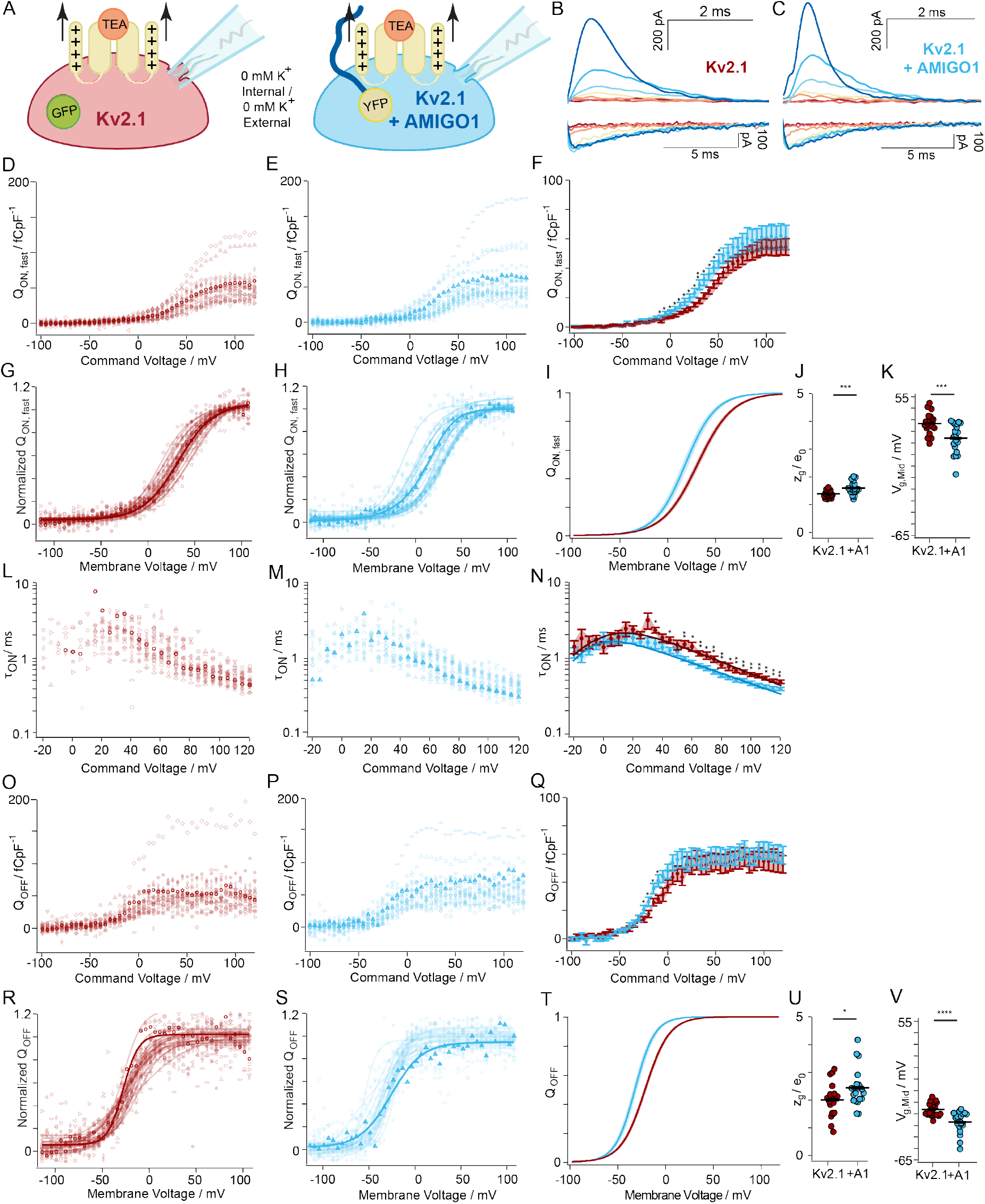
AMIGO1 facilitates the activation of Kv2 voltage sensors. (**A**) Experimental set up: Gating currents (arrows) from Kv2.1–CHO transfected with GFP (red) or AMIGO1–YFP (blue). K^+^ currents were eliminated removal of K^+^ ions and the external tetraethylammonium, a Kv2 pore-blocker (orange). (**B, C**) Top/Bottom: Representative *I*_g,ON_/*I*_g,OFF_ from Kv2.1–control (11.9 pF) or Kv2.1 + AMIGO1 (8.2 pF). Cells were given 100 ms voltage steps ranging from −100 mV (dark red trace) to +120 mV to record *I*_g,ON_ and then stepped to −140 mV to record *I*_g,OFF_. The holding potential was −100 mV. Voltage pulses to −100, −50, −25, +0, +25, +50, and +100 mV are presented. Data points from representative cells are bolded in analysis panels. (**D, E**) *Q*_ON,fast_/*pF–V* relation from individual cells. *Q*_ON,fast_/*pF* is gating charge integrated over the first 3.5 ms normalized to cell capacitance. (**F**) Mean *Q*_ON,fast_/*pF* (**G, H**) *Q*_ON,fast_–*V* relations normalized to maximum *Q*_ON,fast_ from +50 to +100 mV voltage steps. Solid lines represent Boltzmann fit (Eqn. C). (**I**) Reconstructed Boltzmann fits from average *V*_g,Mid,ON,fast_ and *z*_g,ON,fast_ (Table 2). Shading *V*_g,Mid,ON,fast_ ± SEM. (**J**) Steepness and (**K**) midpoint of Boltzmann fits. (**L, M**) *τ*_ON_ from individual cells fit with Eqn. I. (**N**) Average *τ*_ON_–*V*. Solid lines are Eqn. I fit. Fit values ± SD for Kv2.1–control cells: *α*_0mv_ 254 ± 26 s^-1^, *z*_α_ = 0.468 ± 0.026 *e*_0_, *β*_0mV_ − 261 ± 50 s^-1^, *z*_β_ = −1.31 ± 0.37 *e*_0_; for Kv2.1 + AMIGO1 cells: *α*_0mv_ = 443 ± 26 ms^-1^, *z*_α_ = 0.405 ± 0.019 *e*_0_, *β*_0mV_ = 157 ± 52 ms^-1^, *z*_β_ = −2.00 ± 0.55 *e*_0_. (**O, P**) *Q*_OFF_/*pF* relation from individual cells normalized to cell capacitance. (**Q**) *Q*_OFF_/*pF*–*V* relation. (**R, S**) *Q*_OFF_–*V* relations normalized to maximum *Q*_OFF_ from +50 to +100 mV voltage steps. Solid lines are Boltzmann fits (Eqn. C). (**T**) Reconstructed Boltzmann fits using the average *V*_g,Mid,OFF_ and *z*_g,OFF_ (Table 2). Shading *V*_g,Mid,OFF_ ± SEM (**U**) Steepness and (**V**) midpoint of Boltzmann fits. Mean ± SEM. Statistics in Table 2. ****: p = ≤ 0.0001, ***: p = ≤0.001, **: p = ≤0.01, *: p = ≤0.05, ns: not significant. Bars are mean ± SEM.

At voltages above 50 mV, the charge density translocated over the first 3.5 ms, *Q*_ON,fast_, was not significantly different with AMIGO1 (Fig. 6D, E, F), indicating that AMIGO1 did not alter the total charge translocated during early conformational transitions. However, between −10 mV and +50 mV, Kv2.1–control cells did not move as much gating charge as Kv2.1 + AMIGO1 cells, indicating a shift in gating current activation (Fig. 6F). The shift in voltage dependence was quantified by fitting *Q*_ON,fast_–*V* with a Boltzmann (Fig. 6G, H, I) yielding Δ*V*_g,Mid,ON,fast_ of −12.8 ± 3.5 mV (SEM) (Fig. 6K) and a Δ*z*_g,ON,fast_ of 0.215 ± 0.058 *e*_0_ (SEM) (Fig. 6J) (Table 2). This result indicates that AMIGO1 modulates the early gating charge movement which occurs before pore opening.

**Table 2.**
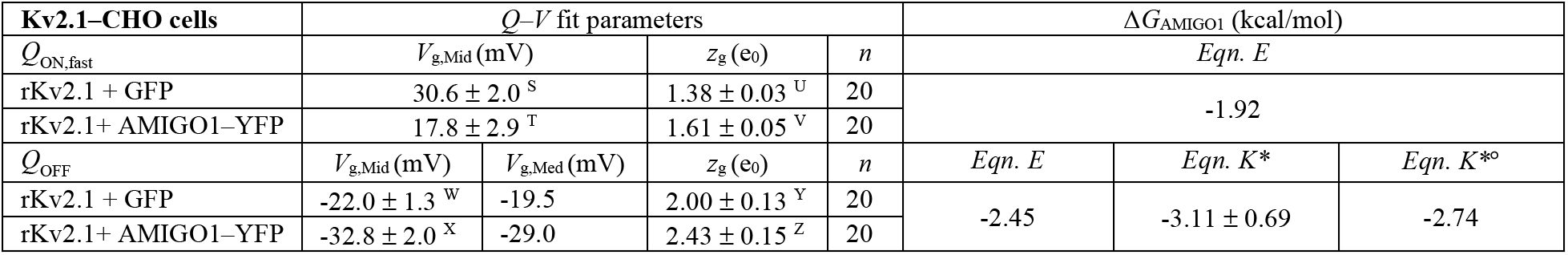
Boltzmann parameters and ΔG calculations for voltage sensor movement. Average *V*_g,Mid_ and *z*_i_ values were derived from 1^st^ order Boltzmann fits of *n* individual cells. Means ± SEM. *V*_g,Mid_ = *V*_g,1/2_. *V*_g,Med_ is median voltage (58). Unpaired, two-tailed t-test p-values: *Q*_ON,fast_: ST: 0.00093. UV: 0.00084. OFF Gating currents: WX: 7.82×10^-5^. YZ: 0.038. \**z* =12.5 *e*_0_, ° *V*_g,Med_ was used.

To determine whether AMIGO1 modulates the kinetics of early gating charge movement, we extracted a time constant (*τ*_ON_) from the decay phase of *I*_g,ON_ that occurs before 10 ms (Fig. 6B top, C top) (Eqn. H) as in (5). In Kv2.1 + AMIGO1 cells, the *τ*_ON_–*V* relation shifts to more negative voltages compared to control (Fig. 6L, M, N). Above +30 mV, the mean *τ*_ON_ for Kv2.1 + AMIGO1 cells was faster than the mean *τ*_ON_ from Kv2.1–control cells (Fig. 6N). Fitting the *τ*_ON_–*V* with rate theory equations indicated AMIGO1 accelerates the forward rate of gating charge movement by 1.7x at neutral voltage and decreases the voltage dependence of this rate by 13% (Fig. 6N). This result indicates that voltage sensors activate faster in the presence of AMIGO1, consistent with destabilization of the earliest resting conformation of the voltage sensors by AMIGO1.

To measure if AMIGO1 alters the total gating charge movement, we integrated OFF gating currents (*I*_g,OFF_) at −140 mV after 100 ms voltage steps (Fig. 6B bottom, C bottom, O, P, Q). The density of *Q*_OFF_ elicited by voltage steps above −10 mV was not significantly different between Kv2.1–control and Kv2.1 + AMIGO1 cells (Fig. 6Q), indicating that AMIGO1 did not alter the density of channels expressed, nor the total gating charge per channel. However, between −25 mV and −10 mV, Kv2.1–control cells did not move as much gating charge as Kv2.1 + AMIGO1 cells, indicating a shift in voltage dependence (Fig. 6Q). Boltzmann fits (Fig. 6R, S, T), yielded Δ*V*_g,Mid,OFF_ of −10.8 ± 2.4 mV (SEM) (Fig. 6V) and a Δ*z*_g,OFF_ of 0.43 ± 0.20 *e*_0_ (SEM) (Fig. 6U) (Table 2), indicating that AMIGO1 shifts total gating charge movement to more negative voltages. Overall, we find that AMIGO1 affects every aspect of gating current we have analyzed to a greater degree than the K^+^ conductance. As both *Q*_ON,fast_–*V* and *α*_0mV_ measurements report the gating charge movements out of the earliest resting conformation, these results indicate that AMIGO1 destabilizes the earliest resting conformation relative to voltage sensor conformations later in the conduction activation pathway of Kv2.1.

### AMIGO1 accelerates voltage-stimulated GxTX–594 dissociation

To further test the hypothesis that AMIGO1 specifically destabilizes the earliest resting conformation of Kv2.1 voltage sensors, we probed the stability of this conformation with GxTX– 594 fluorescence. The earliest resting conformation is stabilized by GxTX (5) and when occupancy of this conformation is decreased by voltage activation, the rate of GxTX–594 dissociation accelerates (45). Destabilization of the earliest resting conformation by AMIGO1 is expected to increase the rate of GxTX–594 dissociation when voltage sensors are partially activated. To test this prediction, we measured the rate of GxTX–594 dissociation at +30 mV, a potential at which about 20% of Kv2.1 gating charge is activated with GxTX bound (5). The rate of GxTX–594 dissociation from Kv2.1 (*k*_ΔF_) accelerated from 0.073 ± 0.010 s^-1^ (SEM) in control cells to 0.115 ± 0.015 s^-1^ (SEM) in cells positive for AMIGO1–YFP fluorescence (Fig. 7). As we see no evidence that AMIGO1 alters GxTX–594 affinity in cells at rest (Supplemental Fig. 5), this 1.6-fold acceleration of *k*_ΔF_ is consistent with AMIGO1 destabilizing the earliest resting conformation of voltage sensors. The thermodynamic model developed to interpret the *k*_ΔF_ of GxTX–594 dissociation (45) estimates that AMIGO1 decreases the stability of the earliest resting conformation of each voltage sensor by 1.9-fold or a Δ*G*_AMIGO1_ of −1.5 kcal/mol for Kv2.1 tetramers (Eqn. L). This result is consistent with AMIGO1 destabilizing the resting voltage sensor conformation to speed up voltage sensor activation and shift conductance to lower voltages.

**Figure 7.**
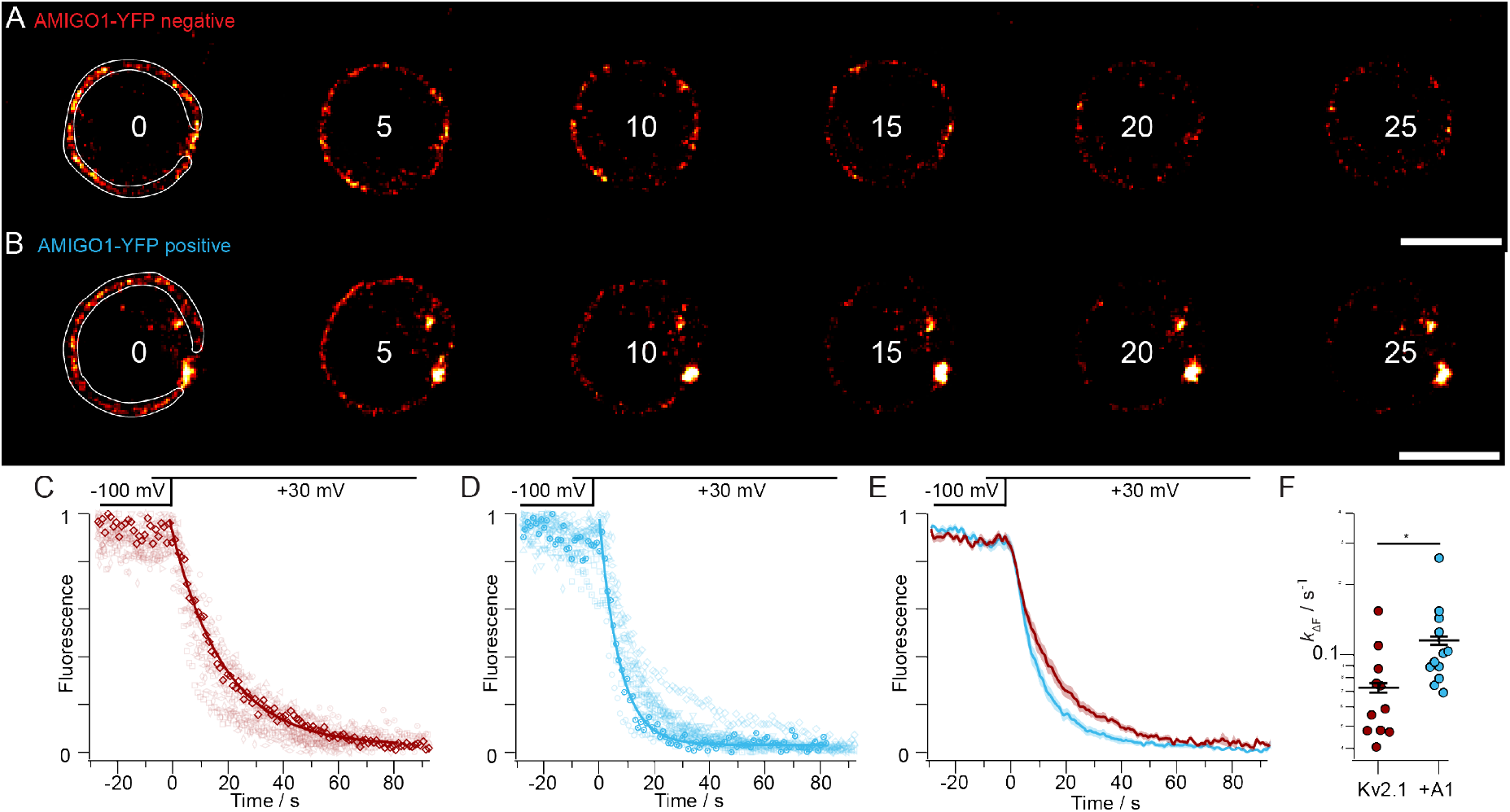
AMIGO1 accelerates voltage-stimulated GxTX–594 dissociation. (**A, B**) Fluorescence from the solution-exposed membrane of voltage-clamped Kv2.1–CHO cells ± AMIGO1–YFP. Kv2.1 expression was achieved through a 48-hour induction period. Cells were held at −100 mV for 30 seconds before being stimulated to +30 mV (time = 0 s) to trigger GxTX–594 dissociation. The time point in seconds of each image is listed. Region of interest for analysis is shown by the white line in left panel, which excludes the point contact with pipette and intracellular regions which have voltage-insensitive fluorescence. 10 *μ*m scale bar. (**C, D**) Normalized fluorescence intensity decay plots for Kv2.1–CHO cells without (red) and with (blue) AMIGO1–YFP fluorescence. The bolded traces correspond to exemplar cells in (A) and (B). Solid line is monoexponential fit (Eqn. G). (**E**) Averaged fluorescence intensity decay for AMIGO1–YFP negative (red), and AMIGO1–YFP positive (blue) cells. SEM is shaded. (**F**) Rates of fluorescence change (*k*_Δf_) were calculated as 1/*τ* from Eqn. G fits. *: *p* = 0.03 unpaired, two-tailed, t-test

### An extracellular surface potential mechanism of AMIGO1 was not detected

To differentiate between mechanisms through which AMIGO1 could change voltage sensor activation we probed whether the large AMIGO1 extracellular domain is directly changing the electrostatic environment of Kv2.1’s voltage sensors. Per surface charge theory, local extracellular negative charges could attract positive gating charges to activate channels (73). AMIGO1 possesses five extracellular glycosylation sites (74), each potentially decorated with negatively-charged sugar moieties (28). AMIGO1 also has a conserved negatively charged residue predicted to be near the extracellular side of the membrane (24, 74). Similar structural characteristics are found in Nav β auxiliary subunits which, like AMIGO1, are glycosylated, single transmembrane pass protein with an immunoglobulin-domain. Nav β1 has been proposed to interact with Nav1.4 α subunit through surface charge effects (75–77). We tested if AMIGO1 likewise affects Kv2.1 activation through electrostatic surface charge interactions.

To measure the electrostatics of the environment immediately surrounding the Kv2.1 voltage sensor domain complex with and without AMIGO1, we employed far-red polarity-sensitive fluorescence (78). The polarity-sensitive fluorophore, JP, was localized to the Kv2.1 voltage sensor by conjugating GxTX to JP at either residue Ser13 or Lys27 (46). When GxTX binds to the extracellular S3b region of the Kv2.1 channel, Ser13 and Lys27 occupy positions of distinct polarity (46). At resting membrane potentials, GxTX Ser13Pra(JP) has an emission maximum of 644 nm, consistent with the homology-based prediction that Ser13 of GxTX localizes in an aqueous environment branched away from S4. Conversely, GxTX Lys27Pra(JP) has an emission maximum of 617 nm, consistent with the prediction that Lys27 sits in the polar region of the membrane adjacent to S4 (46). If AMIGO1 were to alter the electrostatic environment of the resting conformation of the Kv2.1 voltage sensor domain, we would expect either of these environmental point detectors, GxTX Ser13Pra(JP) or GxTX Lys27Pra(JP), to exhibit an altered emission maximum.

Full emission spectra of JP fluorescence from Kv2.1–CHO cells transfected with AMIGO1–YFP and treated with GxTX Ser13Pra(JP) or GxTX Lys27Pra(JP) were fitted with 2-component split pseudo-Voigt functions (Fig. 8C, F). Fitting shows emission peaks, 644 nm and 617 nm, respectively, are unchanged with or without AMIGO1–YFP, consistent with the local electrostatic environment surrounding the JP probes positioned on resting Kv2.1 voltage sensors not being altered by AMIGO1 expression. Previous work has shown that GxTX Lys27Pra(JP) emission peak wavelength is sensitive to conformational changes among early resting states of voltage sensors (46). The absence of any AMIGO1-induced change in environment for either of these GxTX sidechains suggests that AMIGO1 does not cause significant changes to the local environment of the GxTX binding site on the S3b segment of Kv2.1, nor the GxTX position in the membrane when bound to the channel. These results are consistent with destabilization of the GxTX binding site by AMIGO1 being indirect, as the binding site itself appears to retain the same conformation and local environment in the presence of AMIGO1. However, it remains possible that AMIGO1 acts extracellularly to modulate Kv2.1 by a mechanism that these GxTX(JP)-based sensors do not detect.

**Figure 8.**
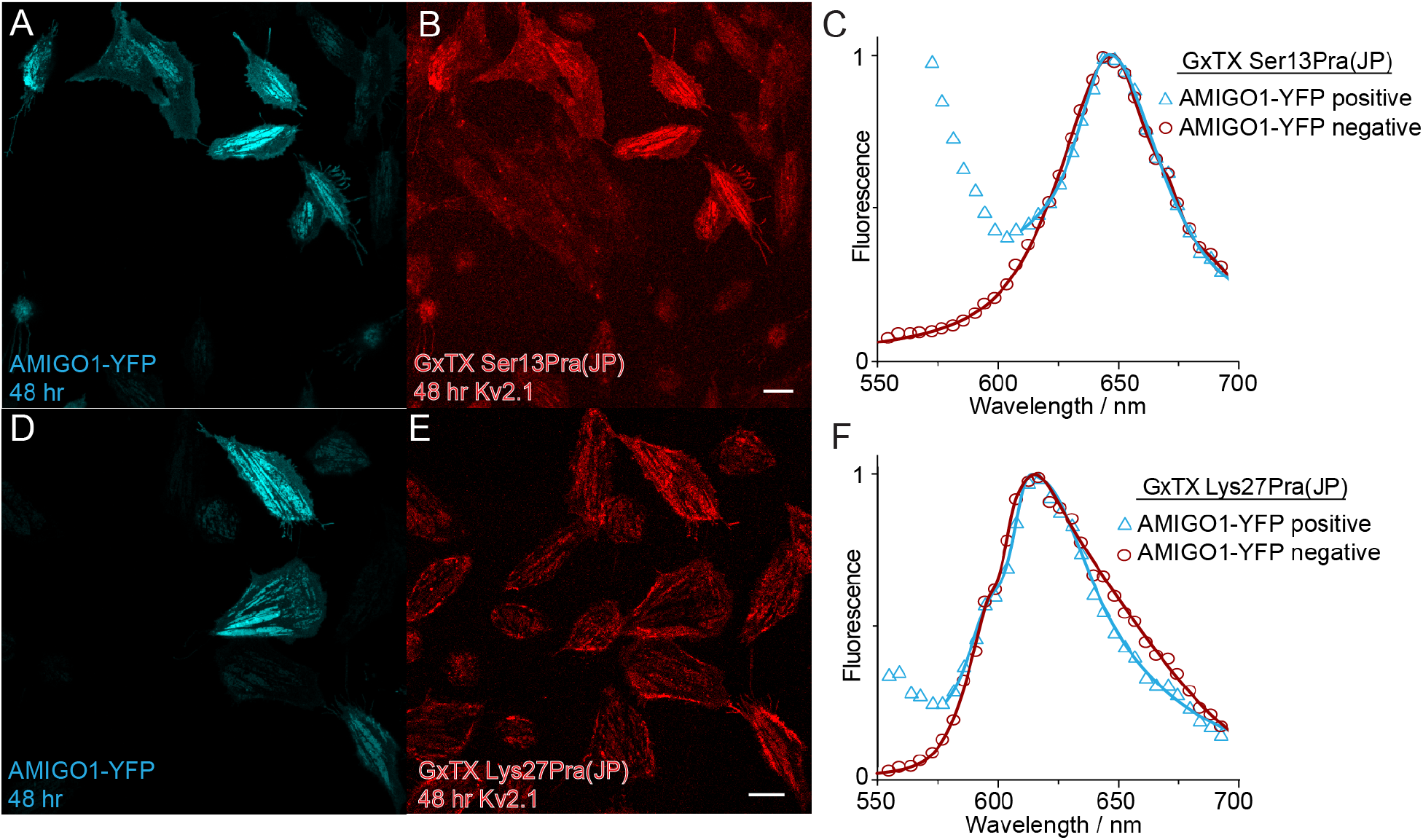
AMIGO1 does not alter the Kv2.1–GxTX interface on resting voltage sensors. Kv2.1–CHO cells transfected with AMIGO1-YFP were treated with GxTX Ser13Pra(JP) or GxTX Lys27Pra(JP) (**A, D**) Confocal image of AMIGO1–YFP fluorescence (blue) and (**B, E**) JP fluorescence. (**C, F**) Fitted emission spectra of cells positive (blue) and negative (red) for AMIGO1–YFP fluorescence. Data points for all spectra are the mean of normalized emission from AMIGO1–YFP positive cells and AMIGO1–negative cells. Spectra were fit with two–component split pseudo–Voigt functions with shape parameters and root–mean–squared values found in Supplemental Table 1.

We also tested whether AMIGO1 acts by a surface charge mechanism with a classical charge screening approach. Surface charge interactions can be revealed by increasing the concentration of Mg^2+^ to screen, or minimize, the impact of fixed negative charges near the voltage sensors (73, 79). If AMIGO1 alters surface potential, we would expect elevated Mg^2+^ to shrink Δ*V*_i,Mid_. To determine whether surface charge screening suppresses the AMIGO1 effect, voltage clamp experiments were conducted as in Fig. 3, except external recording solutions contained 100 mM Mg^2+^ (Fig. 9A, B, C). Kv2.1 requires more positive voltage steps to activate in high Mg^2+^ solutions (Table 1), consistent with sensitivity to surface charge screening (80). In high Mg^2+^, AMIGO1 effected a Δ*V*_i,Mid_ of −7.4 ± 2.4 mV (SEM) (Fig. 10H) but did not change *z*_i_ (Fig. 9G) (Table 1). When compared to low Mg^2+^ conditions by Ordinary 2-way ANOVA, Δ*V*_i,Mid_ was not significantly different in normal versus 100 mM Mg^2+^ (interaction of p = 0.33). Hence, Mg^2+^ altered Kv2.1 activation in a manner consistent with surface charge screening, yet Mg^2+^ did not detectably abrogate the AMIGO1 effect. However, we cannot rule out the possibility of a screened site that is inaccessible to Mg^2+^. While neither extracellular fluorescence measurements nor surface charge screening detected an extracellular impact of AMIGO1, we are not able to rule out the possibility of an extracellular coupling to AMIGO1 that was not detected by these methods.

**Figure 9.**
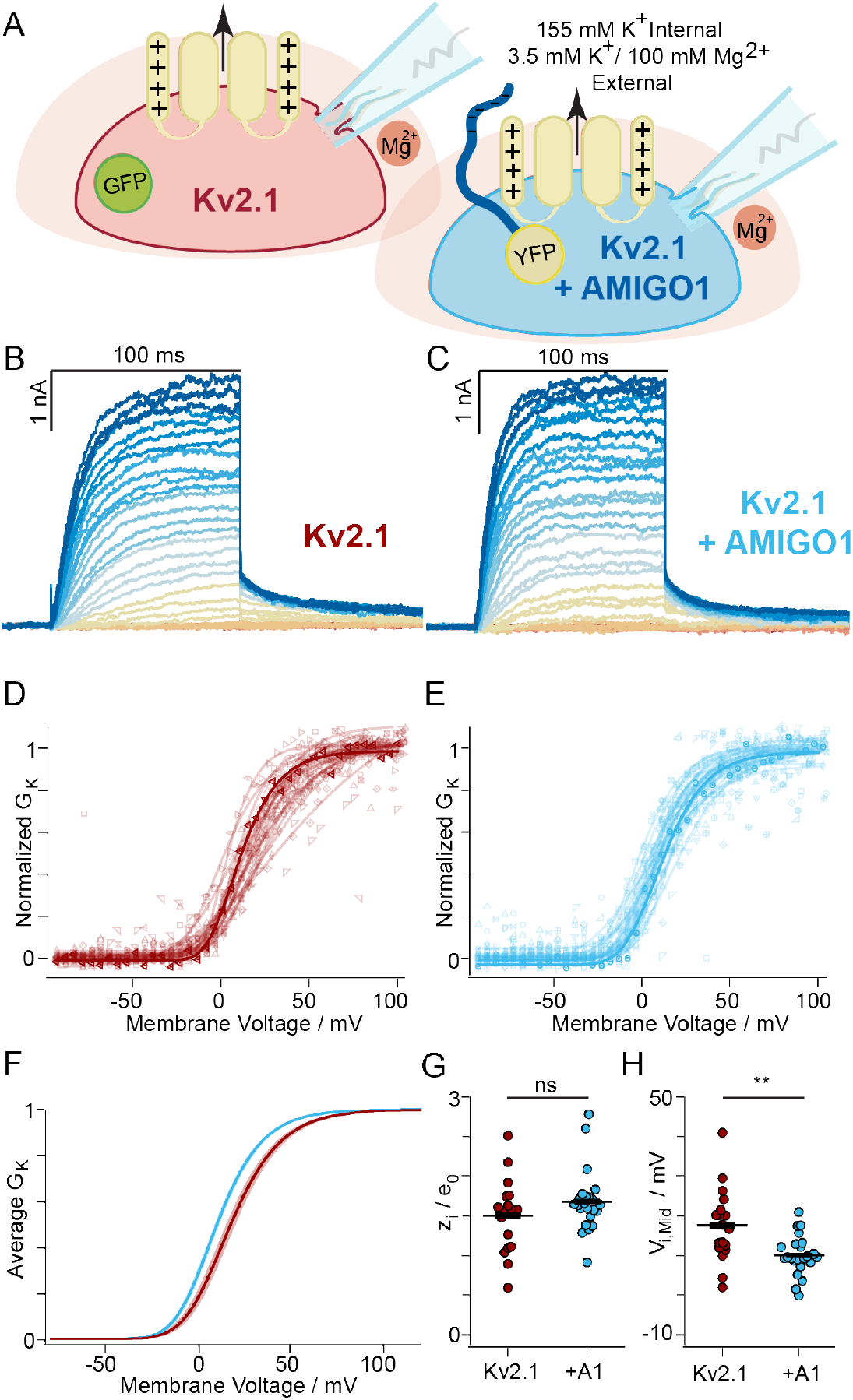
Surface charge screening does not suppress the AMIGO1 effect. (**A**) Experimental set up: Whole-cell K^+^ currents (arrow) from Kv2.1–CHO transfected with GFP (red) or AMIGO1–YFP (blue). 100 mM magnesium was used to shield surface charges (peach halo). Same voltage protocol and representations as Fig. 3. (**B, C**) Representative Kv2.1–control (10.0 pF) or Kv2.1 + AMIGO1 (6.3 pF) cell. (**D, E**) Normalized *G–V* relationships. (**F**) Reconstructed 4^th^ order Boltzmann fits from average *V*_i,Mid_ and *z*_i_ (Table 1). Shading *V*_i,Mid_ ± SEM. (**G**) Steepness and (**H**) midpoint of 4^th^ order Boltzmann fits. Mean ± SEM. Statistics in Table 1. **: *p* = ≤0.01, ns: not significant.

**Figure 10.**
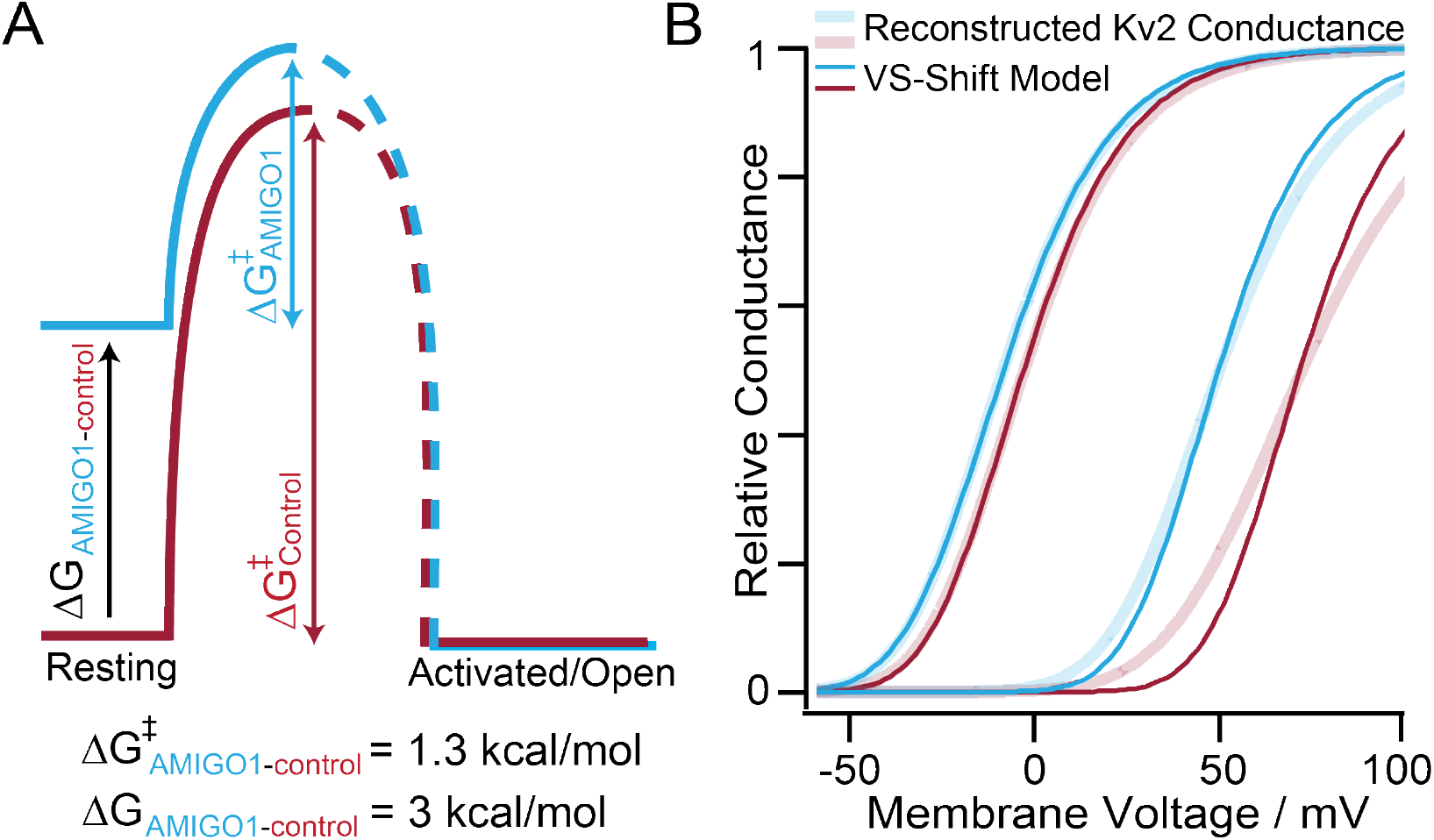
AMIGO1 destabilizes the resting conformation of Kv2.1 voltage sensors. (**A**) AMIGO1 raises resting state energy (Δ*G*) of Kv2.1 voltage sensors and lowers the energy barrier (Δ*G*^‡^) of Kv2.1 activation. (**B**) Voltage sensor shift model of AMIGO1 modulation (dark lines) plotted with reconstructed *G–Vs* from Kv2.1–CHO Table 1 values (pale lines). From left to right: Kv2.1+AMIGO1, Kv2.1–Control, Kv2.1+AMIGO1 with GxTX–594, Kv2.1–Control with GxTX–594. Voltage sensor shift model is 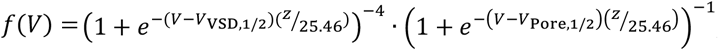, where *z* = 1.5 *e*_0_, *V*_pore,1/2_ = −16 mV, and V_VSD,1/2_ varies. Kv2.1–Control *V*_VSD,1/2_ = −33 mV and Kv2.1–control with GxTX–594 *V*_VSD,1/2_ = 51 mV. AMIGO1 Δ*V*_VSD,1/2_ = −22 mV with or without GxTX–594.

## Discussion

We asked whether AMIGO1 modulates Kv2.1 conductance by modulating conformational changes of pore opening or voltage sensor activation. We found that AMIGO1 destabilizes the resting, inward conformation of Kv2.1 voltage sensors, causing channels to activate at more negative voltages. This conclusion is supported by three major results:

1. *AMIGO1 destabilizes the earliest resting conformation of Kv2.1 voltage sensors*. AMIGO1 expression accelerated conductance activation only at a subset of voltages where the activation kinetics are voltage sensitive (Fig. 3M). When voltage sensor movements were measured directly, gating current recordings revealed an acceleration of the forward rate constant (*τ*_ON_) of gating charge activation in cells with AMIGO1. Between 0 and 120 mV, pore opening is 10-30x slower than *I*_g,ON_ decay (Fig. 3M, 6N), too slow to influence the first few ms of *I*_g,ON_. When the change in the forward rate *α*_0mV_ (Fig. 6N), was used to estimate the amount of energy AMIGO1 contributes to modulating Kv2.1 conformational bias, we found that AMIGO1 imparted −1.3 kcal/mol per channel (Eqn. J) to Δ*G^‡^*_AMIGO1_. From this result we conclude that AMIGO1 speeds the rate of conformational change between the earliest resting conformation and its transition state in the activation path. Additionally, the AMIGO1 effect on GxTX–594 dissociation at +30 mV is consistent with AMIGO1 opposing the action of GxTX–594, which stabilizes resting voltage sensors. All available evidence indicates that AMIGO1 destabilizes the earliest resting conformation of Kv2.1 voltage sensors. We estimate that AMIGO1 destabilizes the fully resting conformation of Kv2.1 channels by ~3 kcal/mol, relative to the fully active open state, and that about half of this energy lowers the barrier for the initial exit of voltage sensors from their resting conformation (Fig. 10A).
2. *AMIGO1 has a greater impact on the voltage sensors than the pore opening*. Free energy estimates indicate more AMIGO1 perturbation of the *Q–V* than the midpoint of the *G–V*. The Δ*G* for AMIGO1’s impact on voltage sensor activation ranged from −1.9 kcal/mol to −3.1 kcal/mol depending on the calculation method (Table 2). Yet, the Δ*G*_AMIGO1_ calculated at the conductance midpoint was only −0.3 kcal/mol (Table 1). This lesser impact on pore opening is consistent with a direct impact of AMIGO1 on voltage sensor movements which are coupled to pore opening. Notably Δ*G*_AMIGO1_ calculated at the conductance midpoint widens to −0.8 kcal/mol when voltage sensor activation is limited with GxTX–594. When we looked at pore opening directly, we saw no evidence suggesting a direct effect of AMIGO1. We saw no change in the slope of the *G–V* relationship with AMIGO1 (Table 1), nor sigmoidicity (Fig. 3), nor single channel measurements (Fig. 4). While these negative results do not eliminate the possibility that AMIGO1 has a small direct effect on pore opening, these negative results constrain the effect size of AMIGO1 on pore opening equilibria to be smaller than the error associated with our measurements.
3. *The AMIGO1 impact on conductance is malleable* In Kv2.1–CHO cells, AMIGO1 shifts the *V*_Mid_ of conductance by −5.7 ± 2.2 mV (SEM). With GxTX–594, the AMIGO1 *G–V* shift widens to −22.3 ± 4.8 (SEM) (Table 1). This remarkable result indicates that the AMIGO1 effect on conductance can change in magnitude. While we have not completely excluded the possibility that AMIGO1 has a direct interaction with GxTX– 594, we think this unlikely, as we saw no sign of an AMIGO1-dependent environmental change around GxTX–JP conjugates, and GxTX–594 had a similar affinity for resting Kv2.1. We think it is more likely that AMIGO1 and GxTX–594 interact only allosterically, and favor the explanation that GxTX makes the *V*_i,Mid_ of conductance more sensitive to the early voltage sensor transition which AMIGO1 modulates. After its fast voltage sensor movement, Kv2.1 has a slow conductance-activating step that makes the 4^th^ power of the *Q–V* not predictive of the *G–V* (3–5, 57). GxTX stabilizes the earliest resting conformation of Kv2.1 voltage sensors such that 4^th^ power Boltzmann fits to the *G–V* are similar to the *Q–V* (5). This suggests the *V*_i,Mid_ is more responsive to AMIGO1 in GxTX–594 because the *G–V* becomes limited by early voltage sensor movement. To test the idea that AMIGO1 modulation of voltage sensors could result in different Δ*V*_i,Mid_ of *G–Vs*, we performed calculations with a voltage sensor shift model composed of simple gating equations. This voltage sensor shift model incorporates distinct *V*_1/2_ values assigned to independent voltage sensor (*V*_VSD,1/2_) and pore (*V*_Pore,1/2_) transitions, all of which must activate to allow channel opening. Calculations incorporating a constant Δ*V*_VSD,1/2_ shift with no change in *V*_Pore,1/2_ demonstrate that the Δ*V*_i,Mid_ of *G–V* can be malleable. In these calculations an AMIGO1 shift of Δ*V*_VSD,1/2_ = −22.4 mV resulted in Δ*V*_i,Mid_ = −5.0 mV (Fig. 10B), similar to the empirical measurement Δ*V*_i,Mid_ = −5.7 mV of Kv2.1 with AMIGO1 (Fig. 3). However, when *V*_VSD,1/2_ was modified to fit GxTX–594 data, this same AMIGO1 shift of Δ*V*_VSD,1/2_ = −22.4 mV yielded a larger shift *G–V* shift, Δ*V*_i,Mid_ = −21.8 mV (Fig. 10B). While the gating model implied by these calculations is highly simplified and does not recapitulate all of our data, it does demonstrate a mechanism by which a fixed modulation of voltage sensors could result in varying Δ*V*_i,Mid_ shifts. As the voltage dependence of Kv2.1 activation is dynamically modulated by many forms of cellular regulation and can vary dramatically (16–20, 81–86), the impact of AMIGO1 might also fluctuate. A malleable impact of AMIGO1 in response to Kv2.1 regulation could perhaps explain why a larger *G–V* shift was originally reported (22), than was observed here or elsewhere (23). The voltage sensor shift mechanism we propose does not require changes in pore opening, or voltage sensor-pore coupling. Maverick and colleagues (23) suggested that the effects of AMIGO proteins on Kv2.1 conductance could be described by increasing the coupling between the voltage sensor and pore opening without a shift in the *Q–V* curve (23), similar to a mechanism by which leucine-rich-repeat-containing protein 26, LRRC26, modulates large-conductance Ca^2+^-activated K^+^ channels (37). As the precise voltage sensor-pore coupling mechanisms for Kv2.1 channels have yet to be defined, we cannot rule out the possibility that AMIGO1 also alters coupling. However, we see no reason that AMIGO1 must do anything other than destabilize the earliest resting conformation of voltage sensors to modulate Kv2.1 conductance.

### Limitations

More detailed investigation of the AMIGO1 impact on the Kv2.1 activation pathway was limited by the relatively small magnitude of AMIGO1-dependent effects versus the cell-to-cell variability, with Δ*V*_i,Mid_ as low as 5 mV, and standard deviations for *V*_i,Mid_ of 4 to 9 mV (Table 1, excluding GxTX–594). While we compensated for the limited power of the AMIGO1 effect by increasing replicates, a decreased cell-to-cell would enable more precise biophysical investigation. This degree of cell-to-cell variability does not appear to be unique to our laboratory. Midpoints reported for rat Kv2.1 activation in HEK293 cells span a 36 mV range, from −20.2 mV to 16.4 mV (22, 23, 67, 72, 87–95). When we calculated *V*_Mid_ standard deviation values from the standard errors and n-values in these studies, standard deviations ranged from 1 to 17 mV, on par with our own. We suspect these notable *V*_Mid_ deviations result from the many different types of regulation to which Kv2.1 channels are susceptible (20, 21). Techniques to constrain the cell-to-cell variability in Kv2.1 function could allow more precise mechanistic studies of AMIGO1 modulation.

Our interpretations assume that the AMIGO1 effect is similar whether Kv2.1 is expressed at low density to measure K^+^ currents or at high density for gating current and imaging experiments. Auxiliary subunit interactions with pore α subunits can be influenced by many factors that can alter their assembly and functional impact on channel currents (96–101). However, if Kv2.1 channels in K^+^ current recording were modulated less by AMIGO1, we would expect a decrease in Boltzmann slope of the fit, a bimodal *G–V* relation, or increased cell-to-cell variability with AMIGO1. We do not observe any of these with CHO cells. The similar impact of AMIGO1 on Kv2.1 conductance in two cell lines (Table 1) and consistency in effect magnitudes with an independent report (23), further suggest that AMIGO1 effect is saturating in our K^+^ conductance measurements. Thus, while incomplete complex assembly and other factors could in theory influence the magnitude of the AMIGO1 impact on Kv2.1 conductance, we do not see evidence that would negate our biophysical assessment of the mechanism through which AMIGO1 alters Kv2.1 conductance.

The most parsimonious explanation for the effect AMIGO1 has on the Kv2.1 conduction–voltage relation seems to be a direct interaction with Kv2.1 voltage sensors. However, it also seems possible that AMIGO1 proteins could change cellular regulation of which in turn modulates Kv2.1. Even if AMIGO1 acts by an indirect mechanism, our mechanistic conclusions remain valid, as they are not predicated on a direct protein–protein interaction between AMIGO1 and Kv2.1.

### Potential physiological consequences of an AMIGO1 gating shift

The impact of AMIGO1 on Kv2.1 voltage sensors suggests that all voltage-dependent Kv2 functions are modulated by AMIGO1. How might the AMIGO1 impact on voltage sensor dynamics affect cellular physiology? As AMIGO1 is colocalized with seemingly all the Kv2 protein in mammalian brain neurons (22, 28, 102), our results suggest that AMIGO1 could cause Kv2 voltage-dependent functions to occur at more negative potentials in neurons. Consistent with this suggestion, *I*_K_ currents from hippocampal pyramidal neurons isolated from AMIGO1 knockout mice are altered compared to wild type *I*_K_ currents (25). AMIGO1 knockout mice display schizophrenia-related features (25) and AMIGO1 knockdown zebrafish have deformed neural tracts (26). However, it is unclear whether these deficits are due to effects on channel gating or other functions of AMIGO1, such as extracellular adhesion. In addition to electrical signaling, Kv2 proteins have important nonconducting functions (28, 67, 103–106), which AMIGO1 could potentially impact. Currently, we can only speculate about whether physiological impacts of AMIGO1 are due to alteration of Kv2-mediated signaling.

Are the AMIGO1 effects on Kv2.1 conductance activation big enough to meaningfully impact cellular electrophysiology? To begin to address this question, we estimated the impact that AMIGO1 would have on neuronal action potentials. In mammalian neurons, Kv2 conductance can speed action potential repolarization (7, 107), dampen the fast afterdepolarization phase (107), deepen trough voltage, and extend after-hyperpolarization (7) to impact repetitive firing (7, 107–110). To estimate the impact AMIGO1 might have on the action potentials, we superimposed the impact of AMIGO1 measured in Kv2.1–CHO cells onto the Kv2 conductance in rat superior cervical ganglion (SCG) neurons, which Liu and Bean (7) found to account for ~55% of outward current during an action potential. We roughly approximated an SCG action potential as a 1.5 ms period at 0 mV, during which the parameters fit by Liu and Bean predict 2.2% of the maximal Kv2 conductance will be activated. If the Kv2 parameters are modified to mimic removal of AMIGO1, SCG neuron Kv2 conductance at the end of the mock action potential decreases by 70% (Table 3). This large effect due to small changes in conductance activation suggests that the AMIGO1 gating shift could have a profound impact on electrical signaling. Furthermore, we think the AMIGO1 impact could be even greater. Liu and Bean found that in SCG neurons, Kv2 activation lacks the slow pore-opening step we see in Kv2.1–CHO cells, and SCG Kv2 kinetics were effectively modeled by a Hodgkin-Huxley n^4^ model of activation (111). This suggests that only voltage sensor activation limits conductance activation in the SCG neurons. When the impact of AMIGO1 on Kv2.1–CHO voltage sensors is applied to SCG neuron parameters, Kv2 conductance at the end of the mock action potential decreases by 89% (Table 3). This analysis suggests that removal of the AMIGO1 effect in neurons could be functionally equivalent to blocking the majority of the Kv2 current during an action potential, which would in turn be expected to have impacts on repetitive firing (7, 107–110). However, we stress that any predicted impact of AMIGO1 on action potentials is merely speculation.

**Table 3.**
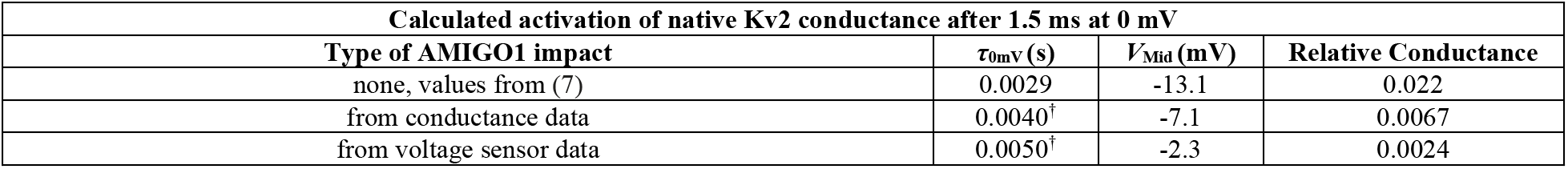
Prediction of AMIGO1 impacts on Kv2 conductance in superior cervical ganglion neurons. Liu and Bean fit Kv2 kinetics with 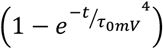 and the *G–V* with 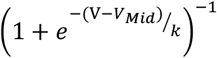, and these equations are used to calculate relative conductance here *τ*_0mV_ and Δ*V*_Mid_ adjusted for the impact of loss of AMIGO1 from Kv2.1–CHO cells. The AMIGO1 impact on conductance activation was a 1.38-fold acceleration of *τ*_0mV_ (Fig. 3M) and *G–V* ΔVi,Mid = −5.7 mV (Table 1). The AMIGO1 impact on voltage sensor activation was a 1.74-fold acceleration of *τ*_0mV_ (change in *α*_0mv_ from fit in Fig. 6N) and *Q*_OFF_–*V* Δ*V*_g,Mid_ = −10.8 mV (Table 2).

### Conclusions

To shift the activation midpoint of Kv2.1 conductance to lower voltages, AMIGO1 destabilizes the earliest resting conformations of Kv2.1 voltage sensors relative to more activated conformations. While we cannot rule out a direct influence on pore dynamics, we saw no indication of such. We propose that AMIGO1 shifts the voltage–dependence of Kv2.1 conduction to more negative voltages by modulating early voltage sensor movements. We also propose that because AMIGO1 acts on early voltage sensor movements, modulation of Kv2 gating can alter the impact of AMIGO1 on K^+^ conductance.

## Author Contributions

Rebecka J. Sepela Conceptualization, Data curation, Formal analysis, Funding acquisition, Investigation, Methodology, Project administration, Visualization, Writing – original draft, Writing – review and editing

Robert G. Stewart

Formal analysis, Visualization

Luis A. Valencia

Formal analysis, Visualization

Parashar Thapa

Resources

Zeming Wang

Resources

Bruce E. Cohen

Conceptualization, Funding acquisition, Project administration, Writing – review and editing

Jon T. Sack

Conceptualization, Funding acquisition, Investigation, Methodology, Project administration, Supervision, Writing – original draft, Writing – review and editing

## Acknowledgements

We thank Vladimir Yarov-Yarovoy, James Trimmer, Karen Zito, and Tsung-Yu Chen of University of California Davis for constructive feedback integral to design and synthesis of experimental ideas. The authors would like to thank the UC Davis statisticians Dr. Sandra Taylor and Dr. Susan Stewart and as well as Karl Brown for their consultation on statistical approaches.

This research was funded by National Institutes of Health grants T32GM007377 (RJS), F31NS108614 (RJS), R01NS096317 (JTS and BEC), and R21EY026449 (JTS). The GxTX-Ser13Cys, GxTX-Ser13Pra, and GxTX-Lys27Pra peptides were synthesized at the Molecular Foundry of the Lawrence Berkeley National Laboratory under U.S. Department of Energy contract DE-AC02-05CH11231. The authors declare no competing financial interests.

---

**Supplemental Figure 1.**
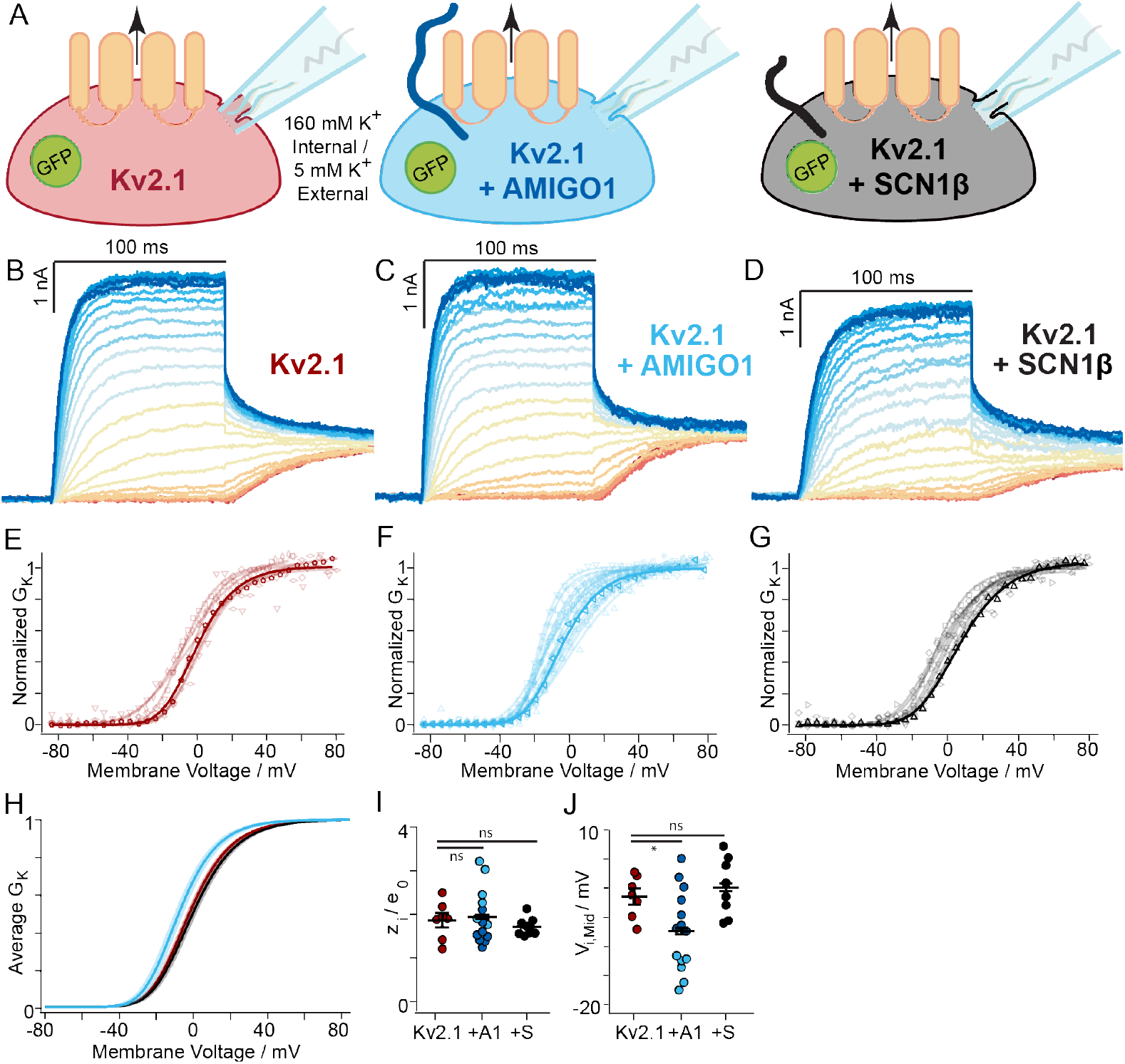
AMIGO1, but not SCN1β, modulates Kv2.1 conductance in HEK293 cells. (**A**) Experimental set up: Whole-cell K^+^ currents from HEK293 cells co-transfected with mKv2.1 and either GFP (red), or AMIGO1–pIRES2–GFP (blue), or SCN1β–pIRES2–GFP (black). (**B**, **C, D**) Representative mKv2.1–control (14.8 pF), mKv2.1 + AMIGO1 (9.6 pF), or mKv2.1 + SCN1β (10.0 pF) HEK293 cell. Data points from representative cells are bolded in analysis panels. (**E, F, G**) Normalized *G–V* relationships for mKv2.1–control, mKv2.1 + AMIGO1, or mKv2.1 + SCN1β cells. Symbols correspond to individual cells. Lines are 4^th^ order Boltzmann relationships (Eqn. C). (**H**) Reconstructed 4^th^ order Boltzmann fits using the average *V*_i,Mid_ and *z*_i_ (Table 1). Shaded areas represent *V*_i,Mid_ ± SEM. (**I**) Steepness and (**J**) midpoint of 4^th^ order Boltzmann fits. For the mKv2.1 + AMIGO1 cells, individual *V*_i,Mid_ and *z*_i_ values are displayed in dark or light blue to highlight an increase in variability. Specifically, the standard deviation of *V*_i,Mid_ increased from ± 3.6 mV in control cells to ± 6.9 mV in mKv2.1 + AMIGO1 cells. We note that the *V*_i,Mid_ values for mKv2.1 + AMIGO1 cells seemed to partition into two groups: a more negatively shifted group with an average *V*_i,Mid_ of −13.9 mV (light blue), and a group similar to mKv2.1 alone with an average *V*_i,Mid_ of −2.5 mV (dark blue). Although all cells analyzed had GFP fluorescence indicating transfection with the AMIGO1–pIRES2–GFP vector, it is possible that some cells were not expressing sufficient AMIGO1 to have a functional effect. Statistics in in Table 1. *: *p* = ≤0.05, ns: not significant. Bars are mean ± SEM.

**Supplemental Figure 2.**
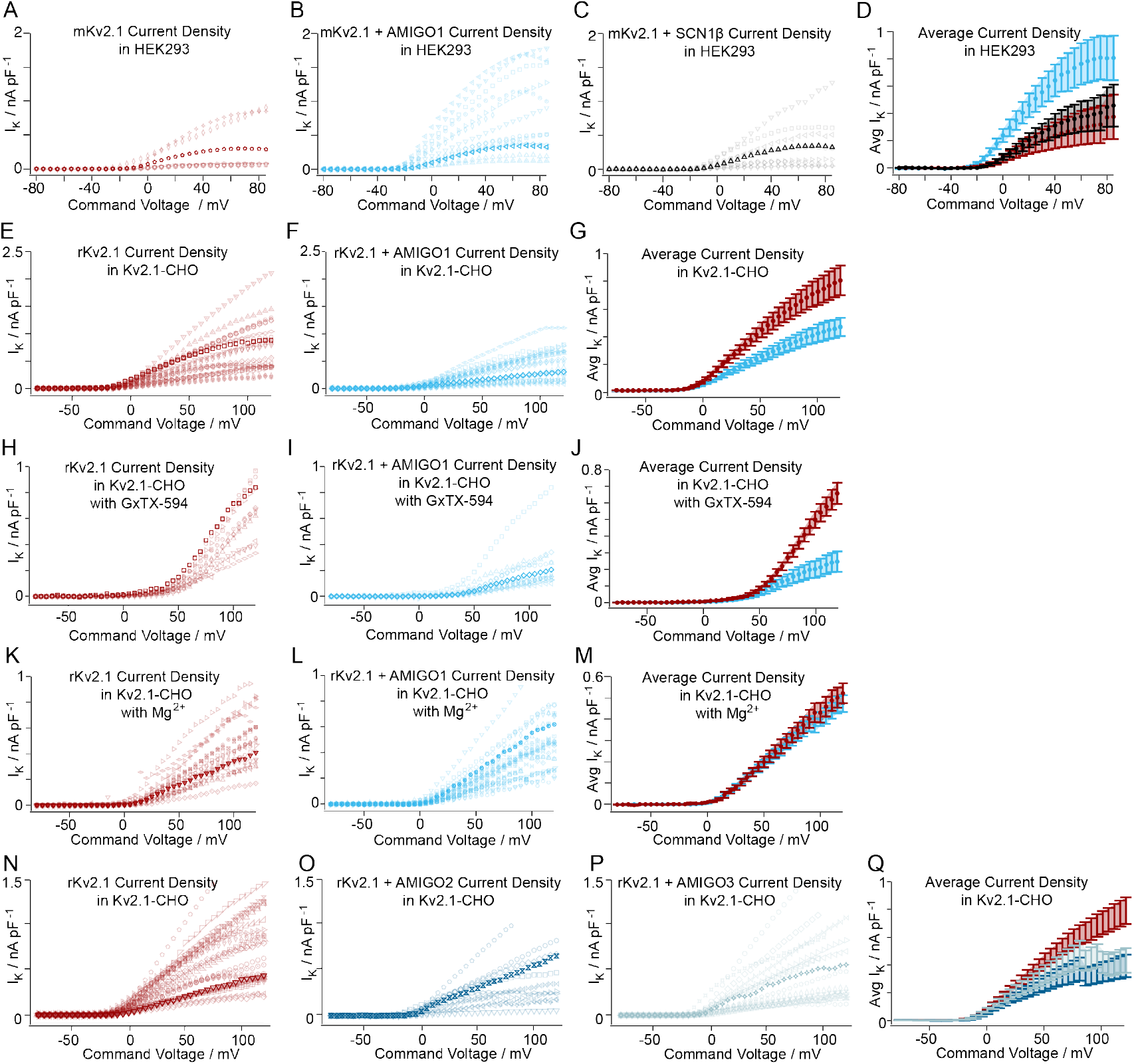
Kv2.1 current density ± AMIGO1 in HEK293 and Kv2.1–CHO cells. AMIGO1 has mixed effects on current density in HEK293 and Kv2.1–CHO cells. Outward current densities normalized by cell capacitance were calculated from mean of the last 10 ms of each voltage step and plotted against the command voltage. Symbols represent individual cells. (**A, B, C**) HEK293 cells co-transfected with mKv2.1 + GFP, mKv2.1 + AMIGO1–pIRES2–GFP, or mKv2.1 + SCN1B–pIRES2–GFP. To limit the proportion of currents from endogenous voltage-dependent channels (53, 60), we set a minimum outward current density as an inclusion threshold for analysis (65 pA/pF at +85 mV). Of the cells patched, 7 of 18 mKv2.1–control cells, 14 of 28 mKv2.1 + AMIGO1 cells, and 8 of 27 mKv2.1 + SCN1β cells satisfied this inclusion threshold and displayed currents consistent with a Kv2.1 delayed rectifier conductance (*I*_K_). Cells that did not meet the inclusion criteria are not plotted making the full variability of current densities is extreme than depicted here. Bolded symbols are exemplars from Supplemental Fig. 1B, C, or D. (**D**) Averages of A, B, and C. (**E, F**) Kv2.1–CHO ± AMIGO1–YFP. Bolded symbols are exemplars from Fig. 3B or 3C. (**G**) Averages of E and F. (**H, I**) Kv2.1–CHO ± AMIGO1–YFP in 100 nM GxTX–594. Bolded symbols are exemplars from Fig. 5B or 5C. Cell symbols matched between E/H and F/I before and after GxTX–594 addition. (**J**) Averages of H and I. (**K, L**) Kv2.1–CHO ± AMIGO1–YFP in 3.5 mM K^+^/100 mM Mg^2+^ external. Bolded symbols are exemplars from Fig. 9B or 9C. (**M**) Averages of E and F. Averaged data are means ± SEM.

**Supplemental Figure 3.**
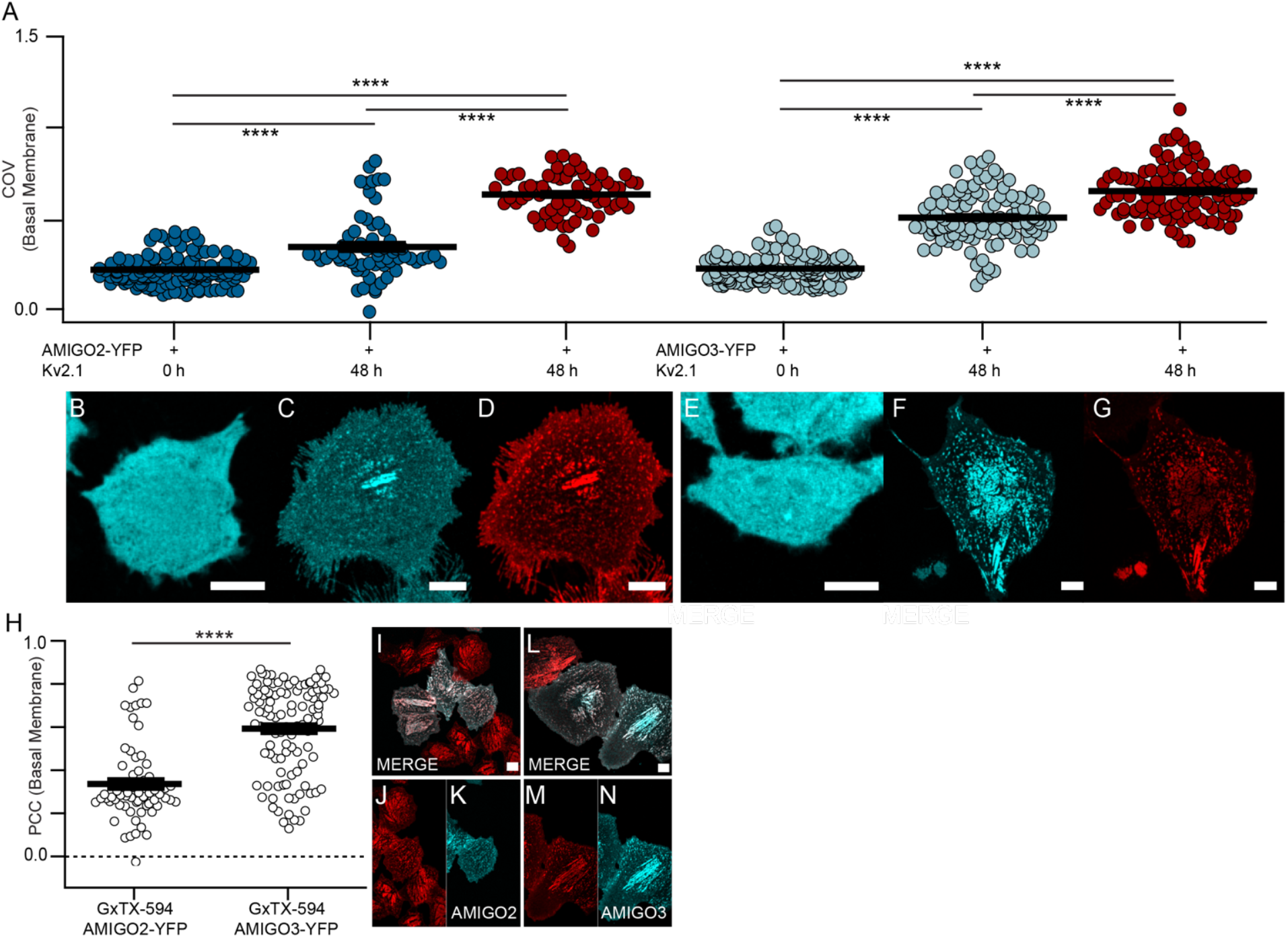
Kv2.1 reorganizes and colocalizes with AMIGO homologs in CHO cells. (**A**) Coefficient of variation of fluorescence from AMIGO2–YFP (dark blue circles), AMIGO3–YFP (light blue circles), or GxTX–594 (red circles). COV from confocal images of glass-adhered membranes (exemplar images in **B-G**). AMIGO2-YFP fluorescence from cells (**B**) not induced for Kv2.1 expression (COVΔ2,oh = 0.2090 ± 0.0062, *n* = 144), (**C**) induced 48 h for Kv2.1 expression (COV_A2,48h_ = 0.342 ± 0.022, *n* = 65). (**D**) GxTX–594 labeling of the cells in C (COV_A2,48h(GxTX–594)_ = 0.631 ± 0.013, *n* = 65 cells). AMIGO3–YFP fluorescence from cells (**E**) not induced for Kv2.1 expression (COV_A3,0h_ = 0.2186 ± 0.0052, *n* = 160), (**F**) induced 48 h for Kv2.1 expression (COV_A3,48h_ = 0.503 ± 0.014, *n* = 109). (**G**) GxTX–594 labeling of the cells in panel F (COV_A3,48h(GxTX–594)_ = 0.650 ± 0.013, *n* = 109 cells). (**H**) Costes thresholded, Pearson’s colocalization coefficients from cells induced for Kv2.1 expression 48 h prior to imaging.. From left to right: PCC_A2,GxTX–594_ = 0.342 ± 0.022, ≥ 0 (p < 0.0001, one–tailed, t-test), *n* = 65; PCC 594 = 0.597 ± 0.020, ≥ 0 (p < 0.0001, one–tailed, t-test), *n* = 108. (**I, J, K)** Exemplar images where merge overlay (white) shows colocalization between GxTX–594 (red) and AMIGO2-YFP (cyan) or **(L, M, N**) AMIGO2-YFP (cyan) Arithmetic means and standard errors are plotted. (**Statistics**) Outliers were removed using ROUT, Q = 1%. An ordinary one-way ANOVA with multiple comparisons was used to evaluate the differences between groups in COV analysis, while a t-test was used to evaluate the PCC data. ****: p = ≤0.0001. Bars are mean ± SEM. All scale bars are 10 μm.

**Supplemental Figure 4.**
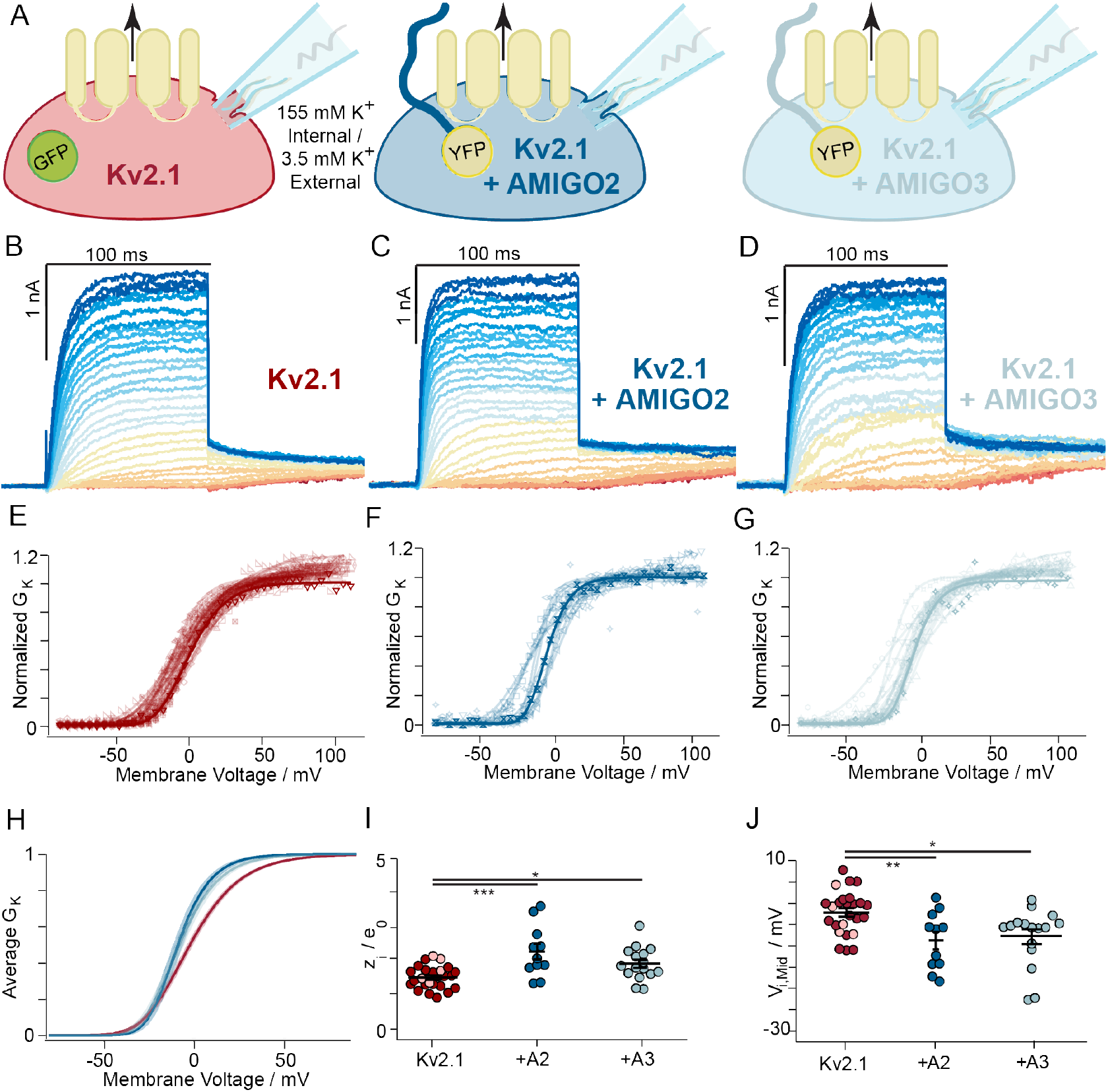
xAMIGO2 and AMIGO3 modulate Kv2.1 conductance in CHO cells. (**A**) Experimental set up: Whole-cell K^+^ currents (arrow) from Kv2.1–CHO transfected with GFP (red), rAMIGO2–YFP (dark blue), or rAMIGO3–YFP (light blue). Same voltage protocols and representation as Fig. 3. (**B**, **C, D**) Representative Kv2.1–control (5.1 pF), Kv2.1 + AMIGO2 (6.6 pF) or Kv2.1 + AMIGO3 (2.4 pF) cells. (**E**, **F, G**) Normalized *G–V* relationships. 5 of the Kv2.1–control cells were recorded from side by side with the Kv2.1 + AMIGO2 cells and Kv2.1 + AMIGO3 cells (light red). There was no statistical difference between these 5 cells and the data previously acquired during Kv2.1 + AMIGO1 recordings for Fig. 3 (assessed by t-test), and data was pooled. Solid lines a 4^th^ order Boltzmann fits (Eqn. C). (**H**) Reconstructed 4^th^ order Boltzmann fits from average *V*_i,Mid_ and *z*_i_ (Supplemental Table 1). Shading *V*_i,Mid_ ± SEM. (**I**) Steepness and (**J**) midpoint of fits. Statistics in Table 1. ***: *p* = ≤0.001, **: *p* = ≤0.01, *: *p* = ≤0.05. Bars are mean ± SEM.

**Supplemental Figure 5.**
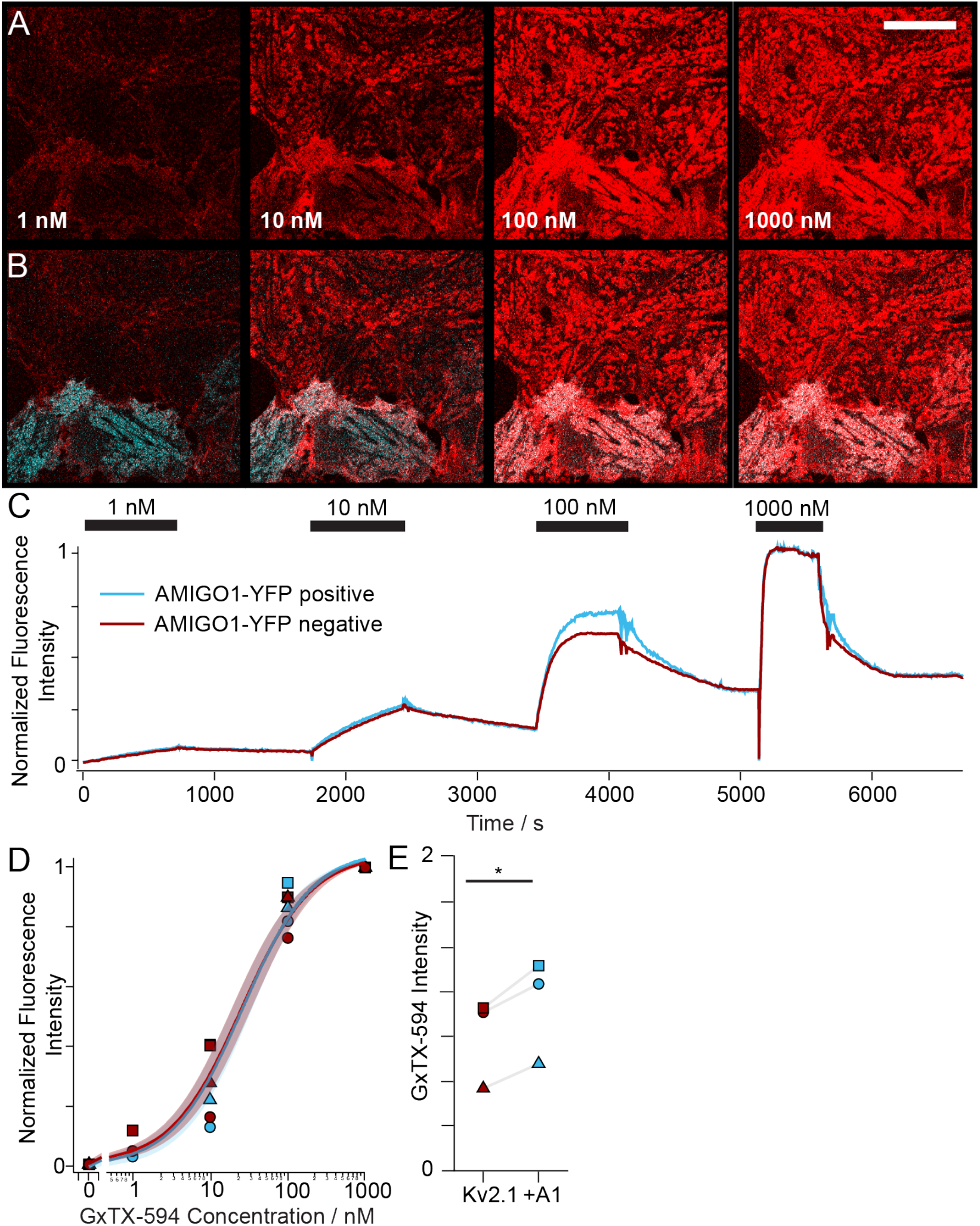
AMIGO1 does not impede GxTX–594 binding to Kv2.1. (**A)** Fluorescence from Kv2.1–CHO cells transfected with AMIGO1–YFP, induced for Kv2.1 expression for 48 hours and labeled with indicated concentrations of GxTX–594 (red). Scale bar 20 μm. (**B**) Overlap (white) between AMIGO1–YFP (cyan) and GxTX–594 fluorescence. (**C**) Mean fluorescence intensities from ROIs encompassing AMIGO1–YFP positive or negative cells from the concentration-response experiment shown in A. (**D**) Normalized fluorescence intensity after 500 s at each concentration as in panel C. Symbol shapes represent data from each of 3 experiments. Curves and shaded regions represent the mean ± SEM of a Langmuir binding isotherm (Eqn. L) fit to individual experiments. *K_d_* = 27.5 ± 8.3 nM without and 27.9 ± 7.2 nM with AMIGO1–YFP. *K_d_* likely is overestimated due to incomplete equilibration at 1 and 10 nM. (**E**) Cells expressing AMIGO1–YFP had brighter GxTX–594 fluorescence with 1000 nM GxTX–594. Symbols correspond with D.

**Supplemental Table 1.**
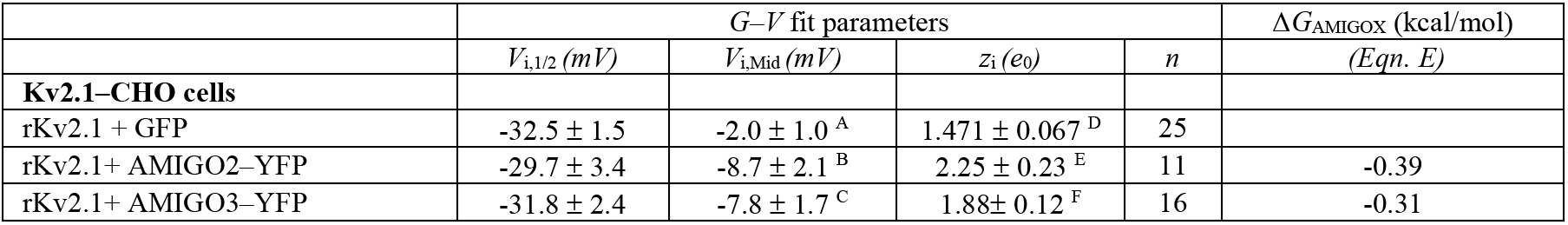
Fourth order Boltzmann parameters for G–V relationships of AMIGO homologs. Average *V*_i,1/2_, *V*_i,Mid_, and *z*_i_ values were derived from a 4^th^ order Boltzmann fits (Eqn. C) of *n* individual cells. All values are given ± SEM. Ordinary one-way ANOVA test with Dunnett’s multiple comparisons p-values: AB: 0.0082. AC: 0.010. DE: 0.0002. DF: 0.026. Δ*G*_AMIGO1_ from Eqn. E, at *V*_i,Mid_ for Kv2.1 + GFP.

**Supplemental Table 2.** Split Pseudo–Voigt fitting parameters. Fluorescence emission spectra split pseudo–Voigt fitting parameters and root-mean squared values.

| GxTX(JP) conjugate | AMIGO1-YFP Expression | fitting component | a0 | a1 | a2 | a3 | a4 | a5 | R<sup>2</sup> |
| --- | --- | --- | --- | --- | --- | --- | --- | --- | --- |
| **GxTX Ser13Pra(JP)** | **- AMIGO** | 1 | 0.229 | 670.4 | 47.88 | 11.41 | 1.075 | 2.323 | 0.999 |
|  |  | 2 | 0.813 | 647.0 | 25.73 | 21.77 | 0.631 | 1.685 |  |
|  | **+ AMIGO** | 1 | 0.893 | 646.7 | 23.30 | 25.63 | 1.822 | 0.721 | 0.997 |
|  |  | 2 | 0.006 | -1610 | -15206 | -1877 | 4967 | 461.2 |  |
| **GxTX Lys27Pra(JP)** | **- AMIGO** | 1 | 0.352 | 594.3 | 12.11 | -11.53 | 0.568 | 5.364 | 0.998 |
|  |  | 2 | 0.719 | 608.2 | 9.71 | 59.05 | 0.359 | -0.264 |  |
|  | **+ AMIGO** | 1 | 0.715 | 597.8 | 16.07 | 18.08 | 1.578 | 2.912 | 0.997 |
|  |  | 2 | 0.632 | 616.3 | 9.05 | 26.28 | -1.657 | 1.488 |  |

**Supplemental Table 3.** N-values for electrophysiology experiments.

| Figure | # Transfections | n per transfection |  |  |
| --- | --- | --- | --- | --- |
| Fig. 3 | 7 | peGFP: 5, 2, 2, 4, 1, 2, 4 | +AMIGO1: 3, 3, 3, 4, 3, 2, 1 |  |
| Fig. 4 | 6 | peGFP: 2, 1, 1, 1, 1, 2 | +AMIGO1: 1, 2, 1, 3, 0, 0 |  |
| Fig. 5 | 5 | peGFP: 4, 2, 2, 3, 2 | +AMIGO1: 3, 3, 1, 3, 2 |  |
| Fig. 6 | 6 | peGFP: 5, 4, 4, 2, 1, 4 | +AMIGO1: 2, 3, 4, 1, 4, 6 |  |
| Fig. 7 | 2 | AMIGO1 (-): 6, 5 | AMIGO1 (+): 5, 6 |  |
| Fig. 9 | 4 | peGFP: 1, 3, 4, 10 | +AMIGO1: 5, 5, 7, 6 |  |
| Sup. Fig. 1 | 4 | peGFP: 3, 3, 1, 0 | +AMIGO1: 4, 4, 6, 0 | +SCNB1: 1, 1, 2, 4 |
| Sup. Fig. 4 | 5 | peGFP: 5, 0, 0, 0, 0<br>(+peGFP n-values from Fig. 3) | +AMIGO2: 1, 2, 0, 1, 7 | +AMIGO3: 1, 7, 5, 0, 3 |

**Supplemental Table 4.** N-values for imaging experiments.

| Figure | # Transfections | # *n* values per transfection |  |  |  |  |  |
| --- | --- | --- | --- | --- | --- | --- | --- |
| Fig. 1 | 4 | YFP (0 hr): 28, 48, 0, 58 | YFP (1.5 hr): 25, 55, 42, 95 | YFP (48 hr): 82, 54, 74, 67 | YFP (ChR): 11, 21, 32, 61 | GxTX-594 (48 hr, AMIGO1): 84, 44, 69, 0 | mRuby-ChR (AMIGO1): 20, 16, 32, 60 |
| Fig. 2 | 4 | AMIGO1-YFP +GxTX-594 (48 hr): 85, 41, 69, 0 |  |  | AMIGO1-YFP +ChR-mRuby: 18, 22, 28, 61 |  |  |
| Fig. 2 | 3 | 0 hr: 41, 35, 25 |  | 1.5 hr: 38, 39, 41 |  | 48 hr: 28, 17, 56 |  |
| Fig. 8 | 3 | AMIGO1(-) (GxTX Ser27Pra-JP): 20, 12, 8 |  |  | AMIGO1(+) (GxTX Ser27Pra-JP): 39, 20, 13 |  |  |
|  | 2 | AMIGO1(-) (GxTX Ser13Pra-JP): 15, 55 |  |  | AMIGO1(+) (GxTX Ser13Pra-JP): 7, 62 |  |  |
| Sup. Fig. 3 | 2 | AMIGO2-YFP (0 hr): 28, 116 | AMIGO2-YFP (48 hr): 59, 6 | GxTX-594 (48 hr, AMIGO2): 59, 6 | AMIGO3-YFP (0 hr): 117, 43 | AMIGO3-YFP (48 hr): 109, 0 | GxTX-594 (48 hr, AMIGO3): 109, 0 |
| Sup. Fig. 3 | 2 | AMIGO2-YFP +GxTX-594: 64, 1 |  |  | AMIGO3-YFP +GxTX-594: 108,0 |  |  |
| Sup. Fig. 5 | 3 | AMIGO1(-): 1, 1, 1 |  |  | AMIGO1(+): 1, 1, 1 |  |  |

